# A Perturb-seq screen guided by species divergence uncovers pathways for collateral artery formation

**DOI:** 10.64898/2026.04.29.721711

**Authors:** Xiaochen Fan, Ronghao Zhou, Brian C. Raftrey, Pamela E. Rios Coronado, Emily Trimm, Erin Clancy, Xinhong Chen, Jamie Bozeman, Maggie S. Chen, Shoxruxxon Alimukhamedov, Juan Alcocer, Idalina Bonham, Stuti Agarwal, Alina Isakova, Vinicio A. de Jesus Perez, Chong Y. Park, Timothy F. Shay, Viviana Gradinaru, Thomas Quertermous, Jesse M. Engreitz, Kristy Red-Horse

## Abstract

Collateral arteries are natural bypasses that can reroute blood flow around arterial blockages, limiting tissue injury during stroke and coronary artery disease. Despite their clinical effectiveness, therapeutic strategies to stimulate collateral artery growth remain unavailable due to our limited understanding of their developmental mechanisms. Remarkably, guinea pigs display exceptionally dense collateral artery networks across various organs, resulting in complete resistance to ischemic damage in the brain and heart. In this study, we compared single-cell RNA sequencing (scRNA-seq) from guinea pig and mouse tissues to identify endothelial cell (EC) gene expression patterns associated with extensive collateral artery development. We then developed an *in vivo* Perturb-seq platform in mice to test whether genes differentially expressed in guinea pigs influence artery EC specification. This pipeline identified artery repressors that were downregulated in guinea pigs and increased pial collateral abundance when inhibited in mice. Downstream analysis suggests that artery repressors, including WNT and hypoxia response genes, function in two capillary EC subsets—*Esm1+* pre-artery and *Apln+* angiogenic tip cells. Reduced activity of these repressors allows more ECs to acquire arterial identity, potentiating collateral artery formation. Collectively, our study establishes a strategy for discovering the genes underlying species-specific traits, suggests that guinea pigs have collaterals due to decreased activity of artery inhibitor pathways and hypoxia responses, and identifies novel targets for stimulating collateral artery formation (**Graphical abstract**).

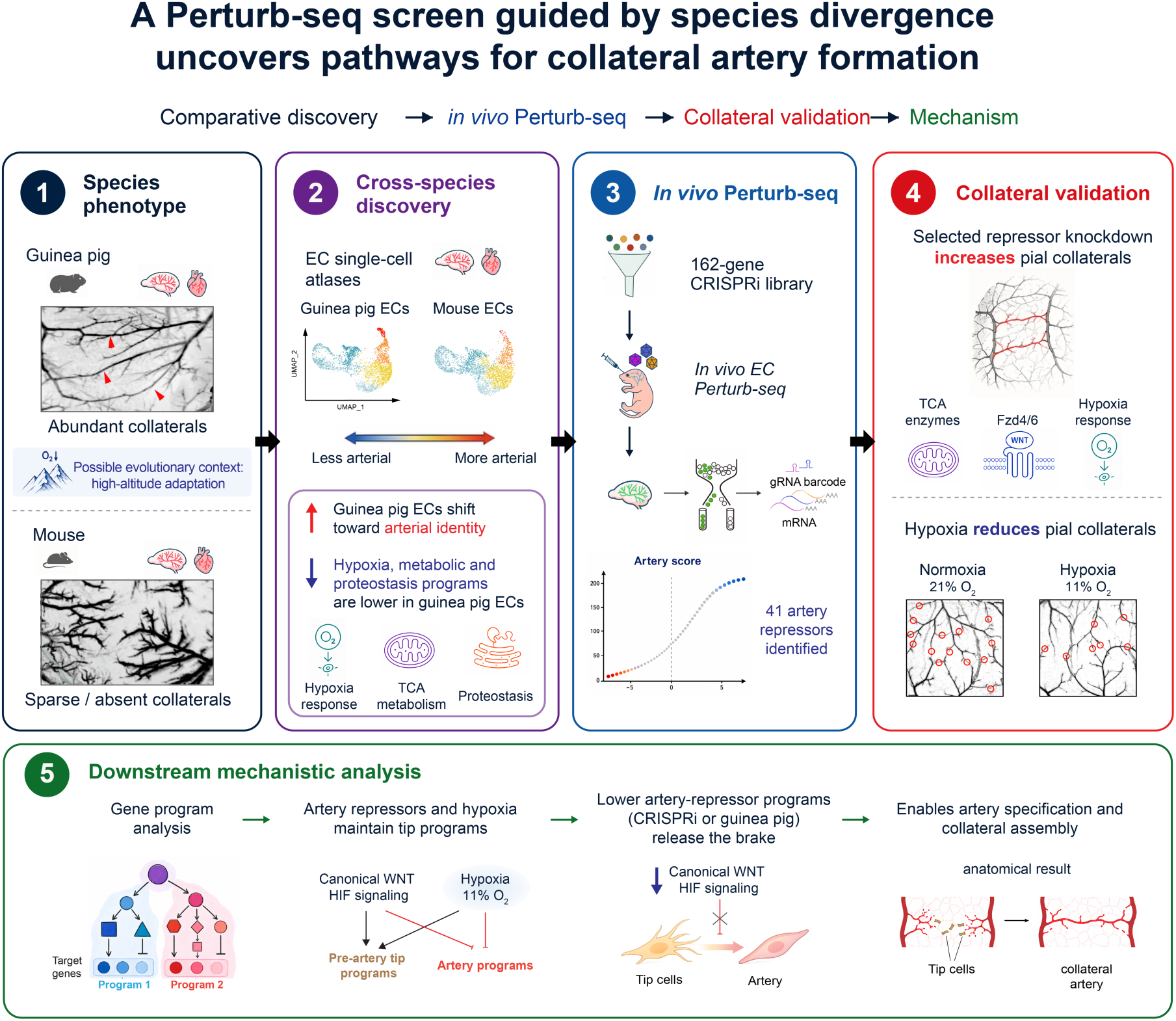

## Introduction

Ischemic vascular diseases such as stroke and coronary artery disease are the leading causes of morbidity and disability worldwide and arise when arterial blood flow becomes blocked ^1,2^. Arterial blockages can be ameliorated by a unique artery subtype called collateral arteries. Collateral arteries form direct connections between two artery branches, like rungs of a ladder, and protect tissues from ischemic damage by providing alternative routes for blood flow ^3–5^. In both the brain and heart, patients with robust collateral networks exhibit significantly smaller infarcts and improved survival rates ^6–10^. Despite their lifesaving capacities, no interventions exist that therapeutically induce collateral formation.

Designing therapies that grow collateral arteries requires an understanding of the mechanisms that govern their formation. To identify these, we turned to an evolutionary outlier: the guinea pig. Unlike all other vertebrate species analyzed, guinea pigs possess dense pial and coronary collateral networks that confer complete protection against ischemic injury ^11–13^. For example, an artery ligation resulting in a large, permanent infarct in the mouse brain or heart barely interrupts blood flow in guinea pigs ^12–14^. Guinea pigs also show increased collateral density in other tissues such as skeletal muscle, indicating that this is a systemic vascular trait, potentially linked to their evolution at high altitudes in the Andes Mountains ^13^. While mice do not naturally form collateral arteries in the heart, they do form these vessels on the pial surface of the brain, the density of which differs among mouse strains ^15,16^. This strain-dependent variation shows that collateral abundance is genetically regulated and suggests that species differences are also genetically encoded. Thus, guinea pigs provide a comparative system to search for molecular pathways that promote collateral abundance, which may provide novel targets for inducing collaterals in other species.

New collateral arteries can form through a variety of cellular processes led by endothelial cells (EC) ^4,17–19^. Artery reassembly occurs when artery ECs migrate out of an existing artery, de-differentiate, proliferate, and coalesce into a connecting branch that differentiates into a mature collateral artery ^20–22^. Arterialization occurs when capillary ECs contribute by undergoing artery specification and either joining an artery EC-derived collateral or forming its own ^22–25^. Regardless of the EC origins, the final maturation step is recruitment of smooth muscle ^21,26,27^. Parts of these complex processes utilize molecular pathways that regulate conventional artery development, such as VEGF, NOTCH, CXCL12, WNT, and hypoxia ^21,28–35^, but there is still much to be learned about the underlying mechanisms, with research efforts in the field focused on the contributions from various EC subpopulations recently identified by single-cell transcriptomics ^19^.

Perturb-seq, which couples pooled CRISPR perturbations with single-cell RNA sequencing (scRNA-seq), provides a scalable framework for linking genetic perturbations to rich transcriptional phenotypes. Because each profiled cell carries both a perturbation identity and a transcriptomic state, this approach can reveal gene functions specific to cell subtypes that may be missed by bulk or low-dimensional assays. Prior Perturb-seq studies have used these rich phenotypes to construct high-dimensional genotype–phenotype maps, infer perturbation-associated pathways, and identify gene function in intact developing tissues *in vivo* ^36–39^.

Building on these frameworks, we combined single-cell comparative genomics with an endothelial *in vivo* Perturb-seq platform to investigate the mechanisms underlying species-divergent collateral formation. We show that guinea pigs form numerous collaterals during embryonic development in both the brain and heart. Cross-species genomics analyses showed that guinea pig ECs are shifted more toward arterial identities when compared to mice and identified candidate pathways differentially expressed in guinea pig ECs. Perturb-seq in the neonatal mouse brain revealed that many of the candidate genes lower in guinea pigs function as repressors of arterial EC specification. Inhibition of a selection of these repressors was sufficient to increase pial collateral formation in mice *in vivo*. Finally, single-cell transcriptional analyses and environmental hypoxia experiments suggest that WNT and HIF signaling function in *Apln+* and *Esm1+* tip cell populations to restrict their arterial specification.

Together, these findings suggest that enhanced collateralization in guinea pigs may arise from reduced activity of artery repressor pathways, promoting a tip-to-artery transition. Our findings also suggest that targeted suppression of these repressor pathways in other species can promote collateral artery formation.

## Results

### Guinea pigs form numerous collateral arteries in the brain and heart during embryonic development

We first determined when collateral arteries form in guinea pigs to identify the developmental window for discovering collateral development genes. Previous studies relied on vascular perfusions to identify artery-artery connections ^11,12^, but this approach is challenging in embryonic tissues and cannot specifically distinguish artery-artery connections from other anastomoses. We therefore tested whether we could identify collaterals using whole organ imaging of immunolabeled artery smooth muscle with anti-smooth muscle actin (SMA) in brains and hearts. With this method, collaterals were readily observed in both organs (**Figure 1a and b**). Mice had collaterals on the pial surface of the brain connecting branches of the anterior cerebral and middle cerebral arteries, but these were much denser in guinea pigs, aligning with previous studies (**Figure 1a**)^3,13,15^. However, species differences were more prominent in the heart—mice had zero collaterals connecting right and left coronary arteries while guinea pigs had a large number throughout the entire volume of the heart (**Figure 1b**). This dramatic difference facilitated quantification in whole organ images, and we therefore carried out a detailed developmental time course in the heart.

**Figure 1.**
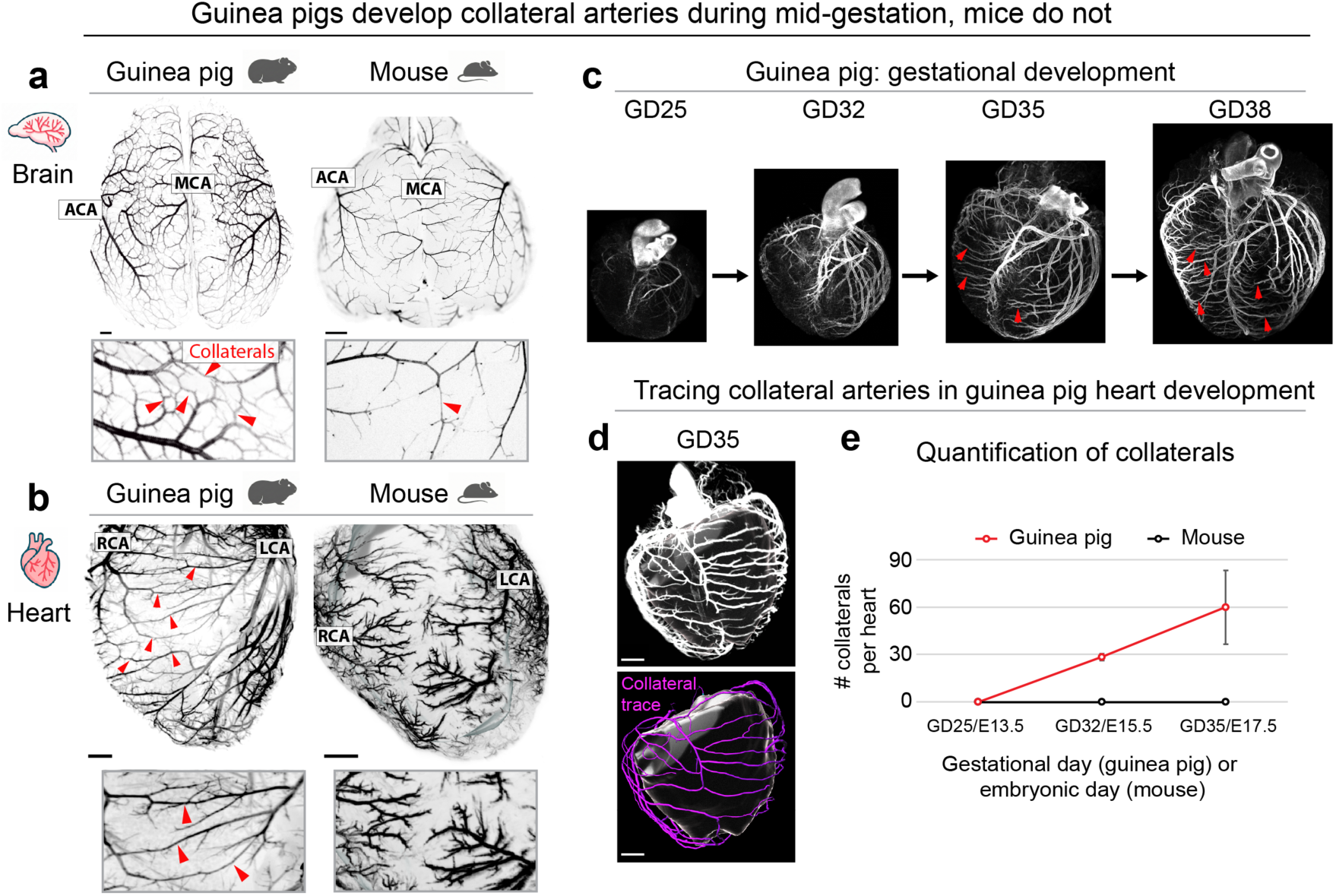
Guinea pigs form abundant collateral arteries during embryonic development. **a and b**, Whole mount smooth muscle actin (SMA) immunostaining (black) to label artery trees in guinea pig and mouse postnatal brains and adult hearts, visualized by light-sheet imaging. **a,** Artery trees in guinea pig and mouse postnatal brains show pial collateral arteries connecting anterior cerebral artery (ACA) and middle cerebral artery (MCA) branches. Insets show higher magnification views; arrowheads indicate representative pial collaterals. Scale bars, 2 mm. **b,** Adult guinea pig hearts possess numerous collateral arteries (arrowheads) connecting the left (LCA) and right (RCA) coronary artery branches. In contrast, mice have none. Insets show higher magnification views. Scale bars, 1 mm. **c and d,** Developmental time course of guinea pig hearts spanning gestational days (GD) 25-38 with whole mount SMA immunostaining (white) to label artery trees and collaterals (arrowheads and purple trace). Scale bars, 250 μm. **e,** Quantification using semi-automated tracing (purple in **d**) showed that coronary collaterals were first evident around GD32 and became progressively more abundant over time (GD25, n=2; GD32, n=2; GD35, n=3).

Guinea pig pregnancy lasts ∼70 days ^40^ with collateral arteries first appearing in embryonic hearts at mid-gestation. Around gestational day (GD) 25, SMA-covered arteries began to form. While these grew over the next few days, they remained as mostly conventional tree-like arteries until approximately GD32-35 when multiple collaterals were evident connecting left and right main coronary branches (**Figure 1c, arrowheads**). Semiautomated tracing (see **Methods**) confirmed that the arteries were collateralized between GD30 and GD35 (**Figure 1d and e**) with an average of 60 connections between the left and right coronary branches at the latter stage (**Figure 1e**). Using a similar embryonic mouse heart whole organ developmental time course as a reference ^41^, we used the onset of coronary artery smooth muscle and extent of branching to match arterialization stages between species (**Table 1**). The stages during which active collateral development occurred were used in the subsequent genomic experiments aimed at identifying guinea pig-enriched collateral effectors.

**Table 1.** Stage matching mouse and guinea pig coronary artery development.

| Coronary artery anatomy | Mouse<sup>41</sup> | Guinea pig (GD) |
| --- | --- | --- |
| No smooth muscle | E11.5-E13.5 | ~GD24-26 |
| Emergence of nascent, tree-like artery branches | E14.5 | ~GD25-30 |
| First appearance of coronary collaterals in GP | Not observed | ~GD32 |
| Mature, highly branched coronary artery tree; numerous collaterals in GP | E16.5+ | ~GD35+ |

### A cross-species, single-cell transcriptomics comparison reveals downregulation of hypoxia, metabolic, and proteostasis pathways

Guinea pigs have higher collateral numbers in the brain, heart, and other tissues ^13^, suggesting the presence of organism-wide EC mechanisms enabling and enhancing collateral development. We hypothesized that differentially expressed genes in ECs from multiple organs would identify drivers of increased collateralization. We performed paired scRNA-seq and ATAC-seq on guinea pig embryonic hearts before (GD25), during (GD32), and after (GD35) collateral development and matched mouse stages (E13.5/15.5/17.5) (see **Table 1**). For the brain, a post-collateral time point was selected for guinea pig (GD35); mouse brain data spanned E18 to P0 because arterial ECs were too rare in the embryonic brain to be reliably captured at a single time point. ECs were enriched by FACS for the brains. Following standard quality control filters (**Figure S1**), the experiment captured >70,000 total cells, with 13,741 brain ECs and 2,422 heart ECs. Joint embedding recovered the expected cell types in both organs from both species (**Figure S2; Figure S3a; Table S1**). We focused this study on ECs for two reasons: prior studies showed that arterial ECs pioneer collateral artery formation ^21,22,25,34^ and increased collateral development occurs across multiple organs in guinea pigs where ECs are the common cellular feature.

In both species, the EC compartments existed as a venous-capillary-arterial continuum (**Figure 2a**), consistent with previous research ^42–44^, and arteries were separated into two clusters: Larger artery (*Gja5*+*Gja4*+*Cxcr4+Aplnr-*) and Arteriole (*Gja5*-*Gja4*+*Cxcr4+Aplnr^low^*) (**Figure 2a-b; Figure S3b**) ^45–47^. Guinea pigs had more ECs in arterial clusters in both organs, increasing by 8% in both the brain and heart (**Figure 2c**). We also calculated artery scores using the top 50 artery marker genes, and these were higher in the total EC population from guinea pigs when compared to mouse (**Figure 2d**). These data suggest that guinea pigs also have a higher proportion of artery ECs in addition to enhanced collateralization.

**Figure 2.**
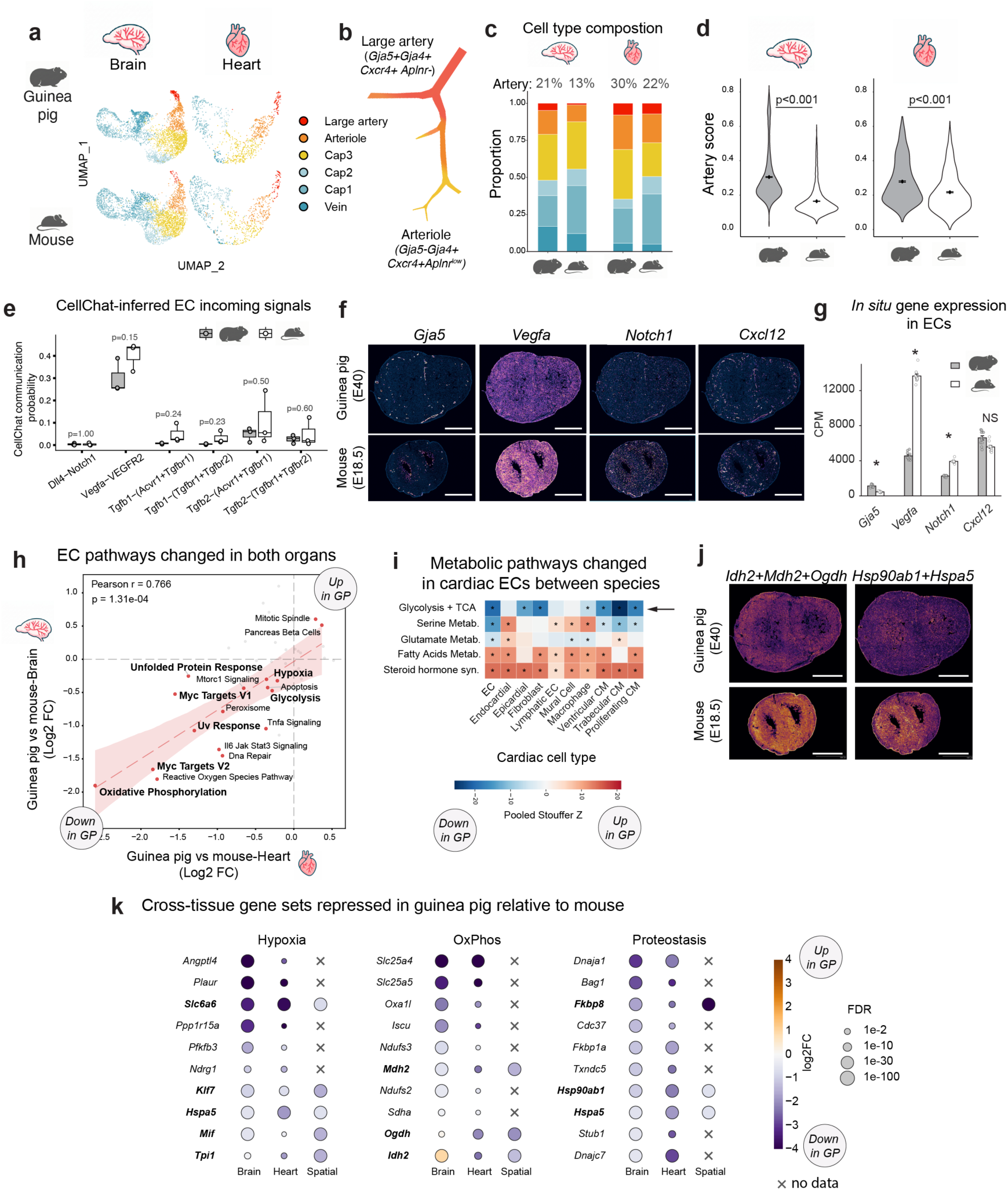
Cross-species genomics comparisons reveal decreased hypoxia and metabolic pathways in guinea pig endothelial cells (EC). **a,** UMAPs of brain and heart EC subsets from developing guinea pig and mouse scRNA-seq, colored by EC subtype. Subsets excluded endocardium for the heart. n=3 samples per species for heart, n=2 samples per species for brain. **b,** Schematic of arterial hierarchy and gene markers used to distinguish the large artery cluster from the arteriole cluster. **c,** Guinea pig brain and heart contain more arterial ECs. Stacked bar graph showing fraction of each EC subtype in brain and heart; values shown are combined proportion of large artery and arteriole. **d,** Overall arterial EC identity is increased in guinea pig. Artery score violin plots for total brain and heart ECs, comparing guinea pig and mouse. p-values indicate between-species comparisons with two-sided Mann-Whitney U tests. **e,** CellChat-inferred incoming signaling to ECs from all cells in heart scRNA-seq does not detect increases in canonical artery development signals (n=3 hearts per species). Communication probabilities shown for indicated ligand–receptor pairs. p-values were generated by two-sided Mann-Whitney U tests. **f and g,** Xenium spatial transcriptomics maps of embryonic hearts. Canonical artery development genes are not up in guinea pig compared to mouse. Scale bars, 1 mm. f, Quantification of EC-specific gene expression is analyzed as pseudobulk across sections with n = 8 sections per heart for each species. Differential expression tested with DESeq2 (negative binomial Wald test, design ∼ section + species) (**g**). **h,** Cross-organ EC pathway changes are concordant and highlight reduced hypoxia, metabolic, and stress-response programs in guinea pig. Each point is a Hallmark pathway plotted as guinea pig versus mouse log2 fold-change (FC). **i,** Metabolic pathway usage is shifted between species, including reduced glycolysis + TCA in guinea pig. Metabolic pathway usage was calculated using single-cell Flux Estimation Analysis (scFEA). **j,** Spatial transcriptomics validates reduced TCA-cycle and chaperone transcripts in guinea pig hearts. Scale bars, 1 mm. **k,** Representative hypoxia, oxidative phosphorylation, and proteostasis genes are consistently repressed in guinea pig across datasets. Dot plots show representative genes from the indicated pathways for ECs from brain and heart scRNA-seq and Xenium spatial pseudobulk gene expression. Color indicates guinea pig versus mouse log2 fold-change, dot size indicates FDR, bold denotes genes measured in all three datasets, and crosses indicate no data.

To identify DEGs that could underlie increased collateralization in guinea pigs, we applied CellChat ^48^ to examine well-established pro-arterial signals, including VEGF, NOTCH, CXCL12, and TGFβ ^21,22,28–31,49–51^. CellChat analyzes gene expression levels and scores curated ligand-receptor interactions based on gene expression per cell type. Heart data were used because they contained all vascular and stromal cell types; brain samples were FACS-enriched for ECs, precluding this analysis. We found that most canonical pro-arterial genes were not higher in guinea pigs, except for an increase in *Tgfbr2* and *Kdr (Vegfr2)* in ECs (**Figure S3c**). However, both *Vegfa* and *Notch1* were increased in the mouse (**Figure S3c**) and no canonical pro-arterial pathway was predicted to be increased in guinea pig ECs by ligand–receptor interaction scoring (**Figure 2e; Figure S3d**). The CXCL12-CXCR4 interaction was not detected because both genes are restricted to a rare artery EC population. This same result was also observed in a heart spatial transcriptomics dataset (stages E18/GD40). Here, while there was an increase in the artery EC identity gene *Gja5* in guinea pig due to the increased number of artery ECs in total, the pro-arterial signaling pathway genes *Vegfa* and *Notch1* were increased in mice (**Figure 2f and g**). These data did not support a model of a direct enhancement of canonical artery specification pathways in guinea pigs; we were therefore motivated to conduct an unbiased search for alternative mechanisms.

We next performed an unbiased analysis of DEGs between guinea pig and mouse total EC populations using GD35/E17.5 brains and hearts. Cross-species differential expression showed strong directionality concordance between the two organs, supporting the hypothesis that organism-wide transcriptomic differences in ECs underlie species-specific vascular development (**Figure S3e**). Comparing gene set activities for the 50 MsigDB Hallmark pathways between species, we found pathways with concordant differences in both organs. Many of the Hallmark pathways were decreased in guinea pig ECs (**Figure 2h; Table S2**). These included hypoxia and metabolic-related programs such as glycolysis, oxidative phosphorylation, and MYC targets as well as genes in the unfolded protein response (UPR) and UV response (**Figure 2h**). A method to infer metabolic flux activity from scRNA-seq data (single-cell Flux Estimation Analysis ^52^; see **Methods**) also predicted reduced glycolysis and TCA cycle activity in guinea pig ECs and, notably, in many of the other cardiac cell types (**Figure 2i**). We validated these gene expression differences with the spatial transcriptomics data, which confirmed the reduction in TCA enzymes (*Idh2*, *Mdh2*, *Ogdh*) and protein chaperones (*Hsp90ab1*, *Hspa5*) (**Figure 2j**). **Figure 2k** shows levels of select individual genes from the Hallmark pathways in **Figure 2h** across all datasets. Based on these data, and emerging research demonstrating the strong effect of metabolic pathways on EC state ^53–56^, we were directed toward a hypothesis in which enhanced collateralization might be a consequence of downregulated hypoxia, metabolic, and/or proteostasis pathways.

### Assessing the effects of guinea pig-enriched and -depleted genes on artery EC specification

Acquisition of artery EC fate is a key component of known collateral development mechanisms ^21,22,25,34^. Prior *in vivo* Perturb-seq studies established that pooled genetic perturbations can be phenotyped across multiple cell types in intact developing tissue by coupling perturbation identity to single-cell transcriptomes ^38,39^. We adapted this logic to test whether species-biased genes regulate arterial specification in a developing vascular bed. This experiment required an *in vivo* model at a developmental stage that was amenable to delivery of pooled guide RNAs and had collateral arteries that could be easily quantified in downstream validations. The neonatal mouse brain met these requirements: brain vascular development extends through early postnatal stages ^22,25,29,30^, brain ECs can be efficiently transduced with EC-tropic AAVs ^57^, and collaterals on the pial surface can be rapidly quantified with standard confocal imaging ^25^.

We designed a CRISPR inhibition (CRISPRi) library of 162 total genes nominated from the cross-species analysis. From the 2,171 DEGs in brain ECs and 2,199 in heart, we selected genes expressed in postnatal ECs of both organs that were likely to act cell-autonomously, and, from this list, further pared down genes using three different analyses: *i)* CellChat: 50 endothelial receptors were selected to represent species-biased signaling inputs; *ii)* CellOracle: 56 transcription factors were selected based on *in silico* predictions of how their deletion would alter arterial fate; and *iii)* Pathway analysis: 40 DEGs were selected based on concordant species-biased expression in both brain and heart, with priority given to hypoxia, metabolic, and proteostasis pathways highlighted by the cross-species analysis. We also included 16 established vascular regulators as benchmarking controls (**Figure 3a, Figure S3f-h; Table S4**). We used a Cre-inducible *CRISPRi* mouse line (*LSL-KRAB-dCas9*) ^58^ crossed with the EC-specific *Tie2Cre* line ^59^ to express CRISPRi specifically in ECs (**Figure S4a and b**). CRISPRi was employed instead of CRISPR knockout in part because it decreases gene expression rather than abolishes it, which likely better approximates the evolutionary expression tuning between guinea pig and mouse and is, therefore, better suited to identify dose-sensitive regulators of arterial fate. A pooled guide library was delivered using the brain EC-tropic AAV9-X1.1 ^57^ at postnatal day (P)1. After incubation for 1 week, perturbation identity was recovered by direct guide capture, adapting direct-capture Perturb-seq strategies in which sgRNAs are sequenced alongside single-cell transcriptomes ^60^ (**Figure 3b**). AAV9-X1.1-GFP showed high infectivity and specificity of ECs as assessed by whole mount immunofluorescence (**Figure 3c**) and flow cytometry (**Figure 3d**), and most cells captured in the scRNA-seq data were ECs (**Figure S4c**).

**Figure 3.**
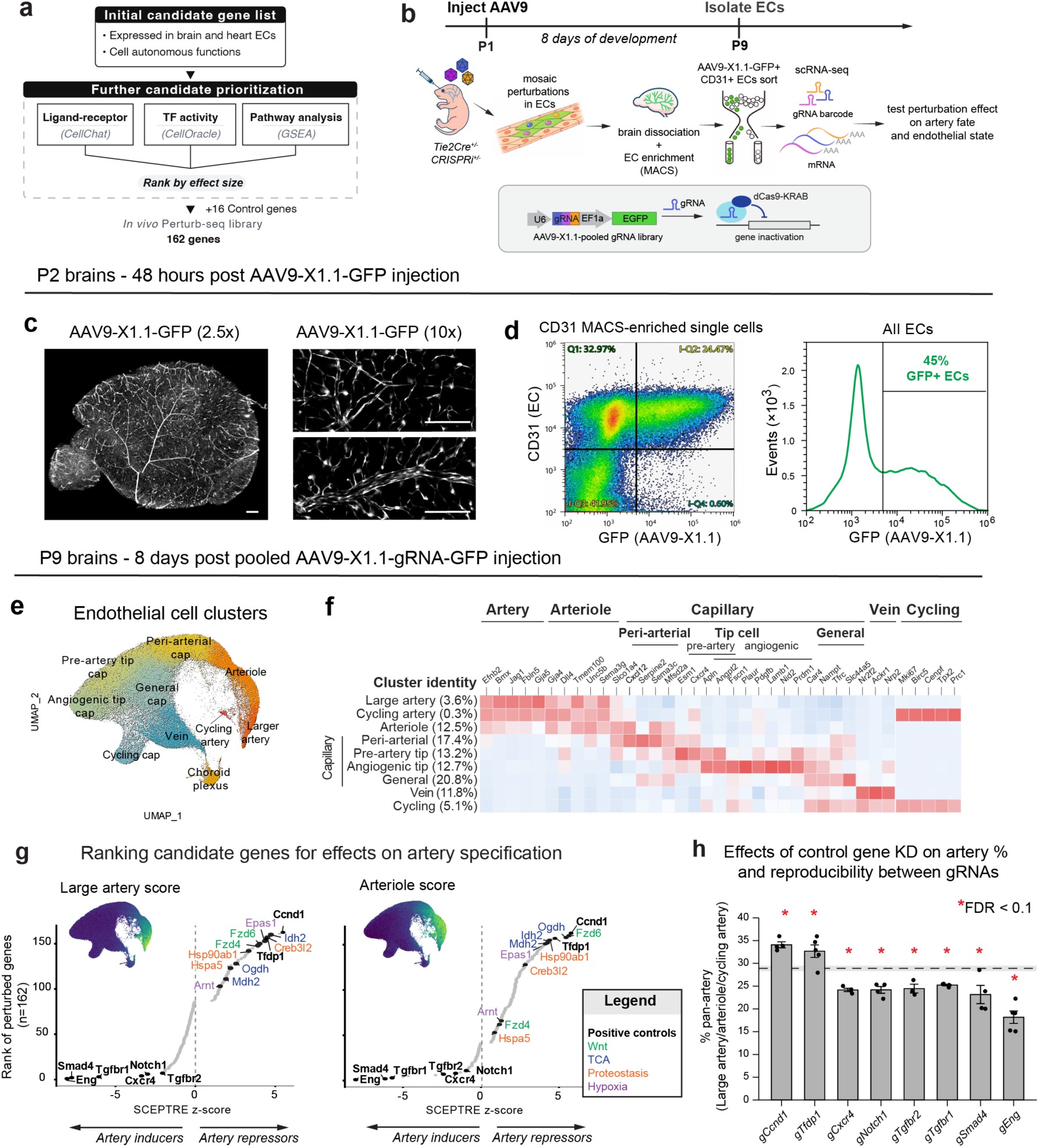
Identification of guinea pig-enriched or depleted artery endothelial cell (EC) fate regulators via *in vivo* Perturb-seq. **a,** Cross-species comparison nominated a 162-gene library for *in vivo* Perturb-seq. **b,** A pooled endothelial CRISPRi workflow enabled screening of artery fate regulators in the neonatal brain. **c,** AAV9-X1.1 broadly transduced neonatal brain vasculature within 48 h. Whole mount confocal images from postnatal day (P) 2 brains after AAV9-X1.1-GFP injection show widespread vascular GFP labeling. Scale bars, 500 μm. **d,** AAV9-X1.1-GFP efficiently labeled brain ECs. Flow cytometry of CD31 MACS-enriched cells and GFP signal across all ECs. **e,** UMAP of ECs recovered from the screen shows that recovered cells spanned the endothelial continuum. **f,** Major EC subsets are annotated based on the expression of the previously studied markers indicated for each EC subtype. Percentages in the dataset are shown. **g,** Ranking perturbation effects recovered known artery inducers and highlighted candidate artery repressors. Rank plots of candidate perturbation effects on large artery and arteriole scores calculated using SCEPTRE. Selected genes are labeled; colors denote pathway classes as indicated. **h**, Control perturbations validated the artery score readout in **g** and demonstrate gRNA reproducibility. Knockdown of known pro-arterial genes decreased the fraction of cells assigned to the artery/arteriole/cycling artery state, whereas knockdown of cell cycle genes increased this fraction. The dashed line marks the control mean, with the gray band indicating its 95% CI. Each dot represents a distinct gRNA targeting the same gene. Bars, mean ± s.e.m. Statistical significance was assessed by Welch’s t-test with Benjamini–Hochberg correction.

EC clusters agreed with published brain and heart atlases ^43,44,47,61–63^ and cluster assignments were guided by prior designations in both organs with one exception: tip-like ECs. The large number of cells, and key information from recent published studies, allowed us to detect two tip cell states—one with an angiogenic-like profile and one similar to the *Esm1+* pre-artery cells described in the retina, intestine, and heart (**Figure 3e and f**, **Table S5**) ^62,64–67^. We denoted three arterial clusters distinguished by connexin expression and cycling genes: “Larger artery” (*Gja5^high^Gja4+*), “Arteriole” (*Gja5^low^Gja4+*), and a rare “Cycling artery” cluster. “Vein” expressed *Nr2f2+*, and a general “Cycling” cluster expressed cell cycle genes. Finally, there were four capillary clusters based on the following markers: 1. “Peri-arterial” representing those adjacent to arteries in the EC continuum with high *Cxcl12* and *Mfsd2a*, 2. “Angiogenic tip” that highly expressed many canonical tip cell markers (*Apln+Angpt2+Pdgfb+Plaur+*) ^63,68–73^; 3. “Pre-artery tip” that expressed tip cell markers at lower levels and the pre-artery tip cell-like markers *Esm1* and *Cxcr4* ^62,64–67^; and 4. a “General capillary” population (**Figure 3f**). The entire EC population also had a distinct choroid plexus EC cluster (*Plvap*+*Mfsd2a-*), but we removed this from our analysis because it has unique properties that were not part of this study (**Table S5**).

The *in vivo* Perturb-seq data passed quality control metrics and displayed proper responses for positive controls. From 10 mice, the data contained ∼260,000 GFP+CD31+ ECs, with a clear barcode rank knee and high library complexity, supporting the quality of the single-cell data (**Figure S4d-f**). 190,300 of the ECs had high-confidence gRNA assignments, and guide multiplicity was centered in the intended ∼5-15 gRNAs per cell range, with robust gRNA UMI detection (**Figure S4g-i**). >90% of targeted genes showed significant mRNA reduction (mean knockdown efficiency ∼40%), and knockdown estimates were concordant between two independent experiments (R^2^ = 0.754; **Figure S4j**). On-target gRNAs produced a much stronger signal for differential gene expression than non-targeting gRNAs, and QQ plots (with SCEPTRE ^74^) showed good calibration (**Figure S4k**). As a biological positive control, we examined *Notch1*, a canonical inducer of arterial fate ^75–78^. Transcriptional changes in *Notch1* knockdown ECs were positively correlated with those induced by *Dll4* knockdown *in vivo* (r = 0.68) ^76^, with concordant down-regulation of arterial markers (e.g., *Efnb2, Cxcr4*) and up-regulation of capillary/tip cell genes (e.g., *Aplnr, Apln, Kdr*) (**Figure S4l**). These results confirm that our screen captures Notch-dependent perturbations to arterial fate consistent with established *in vivo* genetic approaches.

To interrogate the effects of gRNAs on artery specification, we compared the large artery and arteriole gene scores (per cell rank-based signature scores from the top 50 markers) of perturbed cell populations against non-targeting controls using SCEPTRE ^74^ (see **Methods**). Ranking all gRNAs by their SCEPTRE-derived effects revealed genes whose knockdown either increased (i.e. artery repressors) or decreased (i.e. artery inducers) these two artery scores (**Figure 3g**). Importantly, positive and negative controls behaved as expected based on their known functions. For example, the strongest inhibitory effects on artery state were gRNAs targeting four genes in the TGFβ pathway, *Tgfbr1, Tgfbr2, Smad4, and Eng,* which play essential roles in arterial differentiation during early mouse embryogenesis ^50,51,79^ and are key components of our pluripotent stem cell-derived artery differentiation protocol ^80^. Furthermore, ECs carrying gRNAs targeting the canonical arterial activators *Notch1* and *Cxcr4* ^49,75,81^ showed significantly reduced artery and arteriole scores. Conversely, knockdown of the cell cycle regulators, *Ccnd1* and *Tfdp1*, increased arterial identity, consistent with the established requirement for cell cycle exit during arterial differentiation ^42,76,82,83^. These score-based effects were consistent with a secondary analysis assessing effects on the fraction of cells classified as total artery (large artery and arteriole)—for example, TGFβ pathway genes*, Notch1*, and *Cxcr4* knockdowns decreased the arterial EC fraction, whereas *Ccnd1* and *Tfdp1* knockdowns increased it (**Figure 3h**). This analysis also showed that individual guides produced concordant effects (**Figure 3h**).

Beyond the positive controls, most other genes from our candidate list derived from our cross-species analysis behaved in this assay as artery repressors (n = 41), as opposed to inducers (n = 8, FDR < 0.1, **Table S6**). We were particularly intrigued by certain pathways with multiple arterial repressors, which strengthened confidence in the observed effects for individual genes. These included WNT signaling (*Fzd4*, *Fzd6*), TCA cycle enzymes (*Idh2*, *Mdh2*, *Ogdh*), proteostasis (*Hsp90ab1*, *Hspa5*, *Creb3l2*), and the hypoxia response (*Epas1, Arnt, Hif1a*) (**Figure 3g**). These repressor genes were also downregulated in guinea pig ECs from both brain and heart when compared with mice (**Figure 4a**). Thus, our data identified several genes that (i) have lower expression in guinea pigs than in mice and (ii) repress artery EC specification when perturbed *in vivo* in mice, supporting the hypothesis that the decrease in artery repressors in guinea pigs may be related to their increased frequency of collaterals.

**Figure 4.**
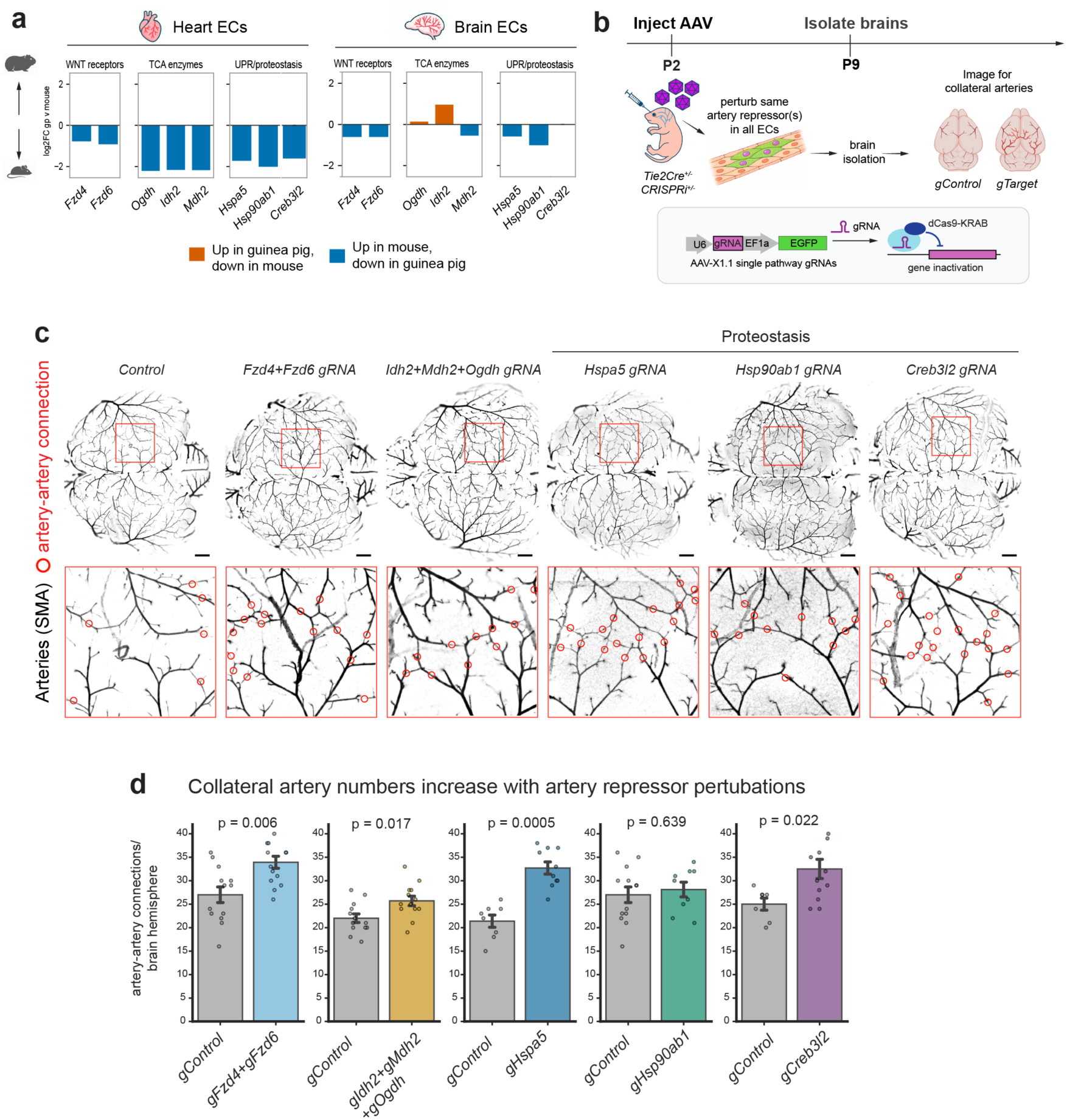
*In vivo* inhibition of artery repressors increases neonatal pial collaterals. **a**, Artery repressors described in Figure 3g are predominantly lower in guinea pig ECs. Cross-species expression differences (log2 fold-change, guinea pig versus mouse) for candidate artery repressors in heart and brain ECs, grouped by pathway. **b**, Neonatal CRISPRi workflow testing artery repressors *in vivo*. **c**, Representative maximum intensity projections of whole brain pial artery immunolabeling using smooth muscle actin (SMA) with zoomed views; artery–artery connections (pial collaterals) are indicated with red circles. **d**, Inhibiting artery repressors increases pial collateral number. Quantification of artery–artery connections per brain hemisphere for each perturbation versus control. For each perturbation, controls are littermates. Dots denote individual hemispheres; bars are mean ± s.e.m. p-values were generated by two-sided Mann-Whitney U tests. Scale bars are 100 μm.

### Inhibiting guinea pig-depleted artery repressor pathways enhances collateral artery formation

The Perturb-seq screen aimed to identify guinea pig-mouse DEGs that regulate arterial EC specification, to nominate candidates that might influence collateral artery formation *in vivo*. Because the candidate selection process (see **Figure 3a**) and screen results (see **Figure 3g**) primarily identified artery repressors that were increased in mice (**Figure 4a**), we were able to utilize the same CRISPRi strategy to inhibit genes in mice and test whether more collaterals form. In this case, gRNAs targeting 1-3 genes in the same pathway were delivered to *Tie2Cre;CRISPRi* pups at P2 via AAV9-X1.1-gRNA-GFPs. Seven days later, at P9, brains were isolated and processed for whole organ confocal imaging and quantification of pial collateral arteries (**Figure 4b**). This experiment tested three of the pathways highlighted in **Figure 3g**: WNT receptors (*Fzd4 + Fzd6*), TCA cycle enzymes (*Idh2 + Mdh2 + Ogdh*), and proteostasis (*Hsp90ab1, Hspa5*, *Creb3l2*).

P9 brains were immunolabeled with anti-SMA to label all arteries, and the entire pial surface was then imaged using confocal microscopy. SMA immunolabeling was used to count the total artery-artery connections for each brain hemisphere. Littermates were used as controls and the researcher was blinded to gRNA identity during counting (**Figure 4c**). We observed significant increases in collateral connections in four of five targets: *Fzd4+Fzd6* (+26%, p = 0.003), *Idh2+Mdh2+Ogdh* (+17%, p = 0.047), *Hspa5* (+53%, p = 0.0005), and *Creb3l2* (+30%, p = 0.007) (**Figure 4d**). Thus, knocking down these genes was sufficient to stimulate collateral artery formation in the neonatal mouse brain (average on-target knockdown in Perturb-seq was 35%).

### Artery repressors influence tip cell and hypoxia gene programs

We next sought to better understand how the artery repressors, many of which were depleted in guinea pig ECs, block artery specification. Using the Perturb-seq data, we identified the specific EC subtypes and gene programs that were affected by perturbing each repressor (**Figure 5a**). This provides insight into how they might restrict artery specification and reveals whether they could work through shared or distinct pathways. Indeed, the artery repressors spanned functionally diverse molecular classes (e.g., WNT receptors, HIF signaling components, protein chaperones) and could therefore affect arteries through different or convergent mechanisms. To investigate this, we used CellRank ^84^ to order the EC subtypes along the vein-artery continuum and interpreted the analyses in the context of this trajectory, as we and others have done in prior scRNA-seq studies (**Figure 5b; Figure S5a-d**) ^42,44,46,67^.

**Figure 5.**
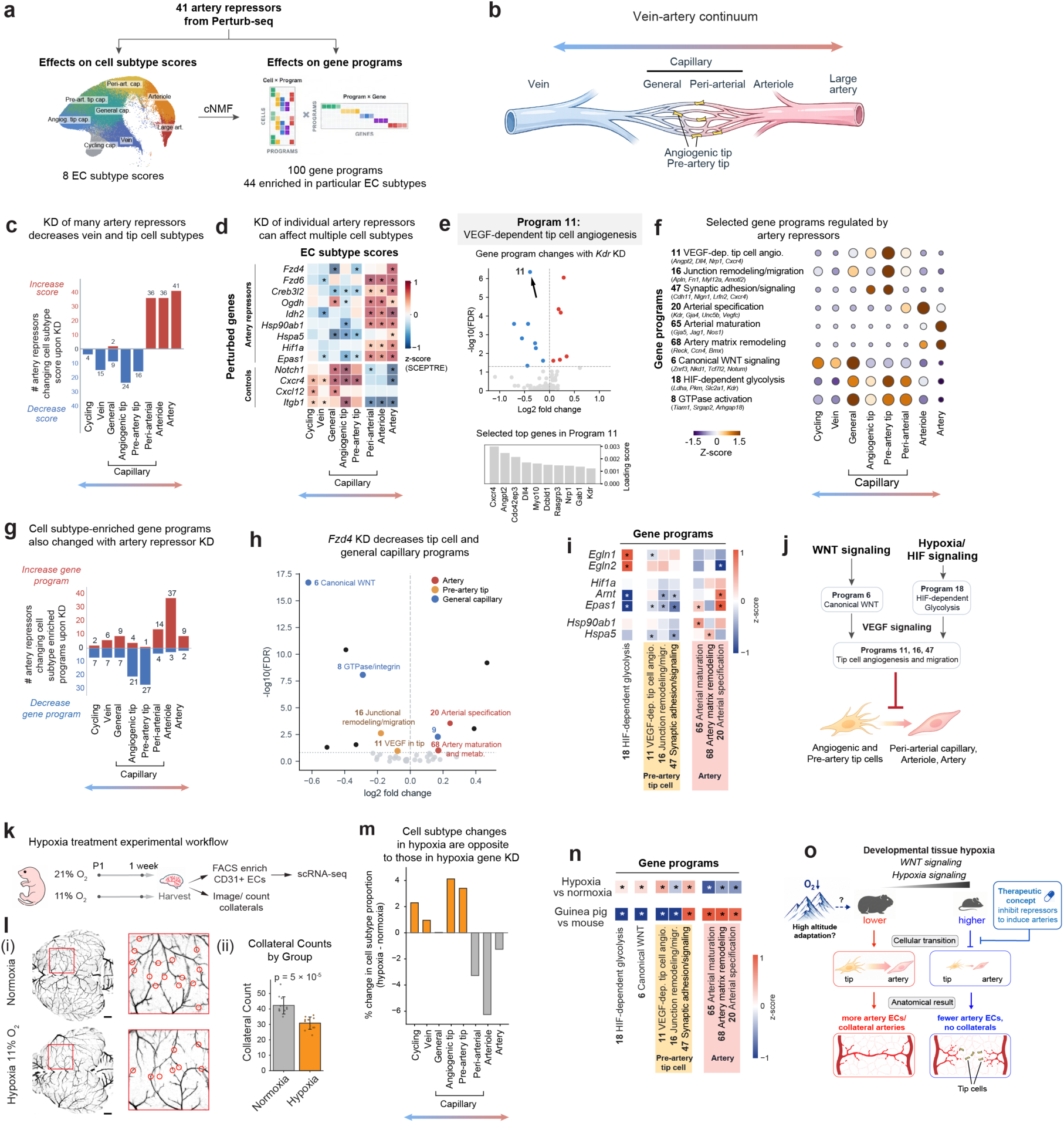
Artery repressors influence tip cells and gene programs downstream of WNT and hypoxia signaling. **a**, Using Perturb-seq data to study how artery repressors affect EC cell subtypes and gene programs. **b**, Localization of EC subtypes along the vein–artery continuum; tip cells are interspersed throughout capillaries. **c**, Artery repressor knockdown preferentially reduces tip cell and vein subtype scores. d, Effects of representative artery repressors on EC cell subtype scores across the vein–artery continuum, shown as SCEPTRE z-scores. Asterisks, FDR < 0.1. **e**, Program 11 represents VEGF-dependent tip cell angiogenesis. **Top**, Program 11 is the most significantly decreased program after *Kdr* knockdown in total ECs; each point is a gene program. **Bottom**, select top genes defining Program 11. **f**, Gene programs are enriched in different EC subtypes. Dot color: mean z-scored program usage. Dot size: fraction of cells with active program usage. Genes listed are selected top-ranked genes for each program. **g**, Tip cell gene programs are the most frequent targets downregulated with artery repressor knockdown. Bars indicate the number of artery repressors whose perturbation increased or decreased usage of enriched programs for each EC subtype. **h**, *Fzd4* loss suppresses pre-artery tip and capillary gene programs while increasing artery programs. Each point is a gene program plotted by log2 fold-change and −log₁₀(FDR); dot color indicates the EC state in which each program is enriched. **i**, Knockdown of HIF pathway and proteostasis genes decreases tip cell gene programs and increases arterial programs. Effects are shown as SCEPTRE z-scores. Asterisks, FDR < 0.15. **j,** Working model whereby genes in the WNT and hypoxia signaling pathways inhibit artery specification. **k**, Neonatal hypoxia experimental workflow. Mice were exposed to normoxia (21% O₂) or mild hypoxia (11% O₂) for 1 week starting at P1, followed by pial imaging and/or FACS isolation of CD31+ ECs for scRNA-seq. **l**, Mild hypoxia reduces pial collateral abundance. (i) Representative smooth muscle actin (SMA)-labeled pial arterial networks under normoxia and 11% O₂; boxed regions are shown at higher magnification, with collateral connections marked with red circles. (ii) Quantification of collateral counts per hemisphere. Bars are mean ± s.e.m. p-values were generated by a two-sided Mann-Whitney U test. Scale bars are 100 μm. **m**, Mild hypoxia expands tip cell subtypes and depletes arterial populations. Changes in EC subtype shown as the proportional difference between hypoxia and normoxia. **n**, Gene programs activated by environmental hypoxia are down in guinea pig. Heat maps show changes in indicated gene programs of interest in hypoxia versus normoxia in mouse ECs and in guinea pig versus mouse ECs under normoxia. Effects are shown as SCEPTRE z-scores. Asterisks, FDR < 0.15. **o,** Working model showing potential interactions between hypoxia pathways and collateral formation in guinea pigs.

We first examined how many of the 41 artery repressors affected each EC subtype to understand whether the increase in arterial cells generally decreased a specific subtype or had wide-ranging effects. Calculating EC subtype scores upon artery repressor knockdown showed that they influenced a variety of EC states, but that some were more affected than others (**Figure 5c**). Angiogenic tip cells were the most affected with their cell type score decreasing with knockdown of 24 of the 41 repressors. Vein and pre-artery tip cells followed close behind with 16 and 15 repressors, respectively. Interestingly, knockdown of most artery repressors not only increased arteriole and large artery scores, but also peri-arterial capillary scores (**Figure 5c**). Individual repressors often reduced multiple cell states; for example, perturbing the WNT receptor *Fzd4* decreased EC subtype scores for both general capillary and pre-artery tip cells while *Epas1* decreased vein and both tip cell subtypes (**Figure 5d**). These data show that, while individual repressors act on different EC subtypes, there is a bias toward affecting the two tip cell subtypes with an inflection point in the vein-artery continuum between pre-artery tip and peri-arterial capillary cells.

Because perturbation effects can be distributed across coordinated transcriptional programs rather than captured by individual marker genes alone, we next summarized the screen in a Perturb-seq-derived gene-program space ^36–38^. We applied consensus non-negative matrix factorization (cNMF ^85^) to decompose the Perturb-seq data into 100 co-regulated gene programs—i.e., genes that are co-expressed across single cells and perturbations (**Figure S6a-c**). We also tested which gene perturbations regulated the expression of each gene program using SCEPTRE ^74^ (**Figure S6d; Table S8**). Together, this information was used to annotate each program based on: *i)* top-loading genes, *ii)* EC subtype enrichment, and *iii)* the identity of program regulators (**Table S9**). For example, Program 11 was annotated “VEGF-dependent tip cell angiogenesis” because its top-loading genes were established components of the VEGF–tip cell axis (*Cxcr4*, *Angpt2*, and *Nrp1*) ^69–73^, it was enriched in the tip cell states, and it was the top program downregulated upon *Kdr* perturbation (**Figure 5e and f**). Other example gene programs are listed in **Figure 5f** and further explained in **Figures S6e-i** and **Table S9**. Interactive visualizations of program annotations and perturbation effects on all gene programs are available at the <u>manuscript resources page</u>.

We next compiled 44 gene programs significantly enriched in individual EC subtypes (each subtype had between 3-8 enriched programs, which sometimes overlapped) and asked which of these programs were also changed by knockdown of the 41 artery repressors. Again, tip cell subtypes were prominently involved; among the 44 subtype-enriched gene programs, artery repressor knockdown decreased tip cell-enriched gene programs more than any others (21 artery repressors affected programs in angiogenic tip and 27 did so in pre-artery tip) (**Figure 5g**). As for specific programs contributing to enrichment in tip cells, the highest-ranking programs in terms of how many repressors decreased each program were the three programs enriched in pre-artery tip cells (Programs 16, 47, and 11), which were decreased with knockdown of repressors such as *Fzd4, Fzd6, Ogdh, Hspa5,* and *Kdr* (**Figure S6j and k**).

We then examined select artery repressors and their potential effects in tip cells. First, we considered the WNT receptor *Fzd4* as an example. Consistent with the role of *Fzd4* as an endothelial Frizzled receptor linked to canonical WNT–β-catenin signaling ^86,87^, *Fzd4* knockdown decreased Program 6–Canonical WNT signaling (**Figure 5h, Figure S6h and j**). *Fzd4* knockdown also affected the expression of 11 other programs, which were distributed across cell subtypes (**Figure 5h**). Pre-artery tip cell-enriched Program 11-VEGF-dependent tip cell angiogenesis and Program 16-Junctional remodeling and migration were down, and general capillary Program 8-GTPase-Integrin Cytoskeletal Signaling was also down (**Figure 5h**). Considering the effects on both EC subtypes (**Figure 5d**) and gene programs (**Figure 5h**), these data suggest that *Fzd4*-mediated canonical WNT signaling may suppress artery specification in part by promoting VEGF-dependent and migration gene programs in pre-artery tip cells.

Finally, we considered our unexpected finding that genes in the hypoxia response pathway can repress artery specification in ECs. We examined the canonical HIF transcriptional components (*Hif1a, Epas1, Arnt*) and the HIF-destabilizing prolyl hydroxylases *Egln1* and *Egln2* ^88–90^, alongside the proteostasis chaperones *Hsp90ab1* and *Hspa5*, which stabilize HIF protein and thereby enhance hypoxia signaling ^91,92^. Under hypoxia, the EC-enriched dimer EPAS1/ARNT (also known as HIF-2α/HIF-1β) transcriptionally activates *Vegfa* and glycolytic regulators ^93,94^, a response suppressed by EGLN1/2 in normoxia ^88–90^. Consistent with this pathway architecture, *Arnt* and *Epas1* knockdown suppressed Program 18-HIF-dependent glycolysis, while *Egln1* or *Egln2* knockdown induced it (**Figure 5i; Figure S6i and j**). *Hif1a* knockdown was directionally similar to *Arnt* and *Epas1* but did not reach significance. Thus, perturbing HIF pathway genes has the expected effect on the hypoxia-induced glycolysis gene program.

What specific EC subtypes and gene programs did hypoxia pathway members affect? Tip cells were the most decreased subtype among the different hypoxia-related perturbations. *Epas1* and *Hspa5* knockdown decreased both tip cell subtypes while *Hsp90ab1* knockdown decreased angiogenic tip cells (**Figure 5d**). *Arnt, Epas1, and Hspa5* knockdown also decreased one or more of the three pre-artery tip cell-enriched programs (Programs 11, 16, and 47), while increasing artery programs (Programs 65, 68, and 20) (**Figure 5i**). Although not significant, *Egln1/2* trended in the opposite direction for multiple tip cell programs (**Figure 5i**). These results suggest that hypoxia response genes may inhibit artery specification at least partly through gene programs in tip cell subtypes.

Together, these data support a model in which multiple artery repressors, including genes in the WNT and HIF signaling pathways, restrain artery development by inhibiting capillary-to-artery specification, particularly in tip cell subtypes through tip cell-enriched gene programs (**Figure 5j**).

### Environmental hypoxia affects tip cell programs and suppresses collateral formation

We sought to use orthogonal methods to validate our unexpected finding that hypoxia genes in ECs inhibit artery specification because: 1) genes in this pathway had consistent effects on EC programs and artery specification (**Figure 5d and i**); 2) oxygen levels are an environmental factor that differed during life histories between mice (evolved at low elevation) and guinea pigs (evolved at high elevation) ^13^; and 3) mutations that decrease hypoxia responsiveness are the most consistent genetic variants implicated in evolutionary adaptation to high altitudes in other organisms, including human populations ^95–97^.

To test whether environmental oxygen levels would influence collateral artery formation, or the aforementioned artery and tip cell programs, we exposed mouse pups to normoxia or mild hypoxia (11% O₂) for one week starting at P1, followed by pial artery imaging with blinded collateral counting or FACS isolation of brain ECs for scRNA-seq (**Figure 5k**). Consistent with the gene-level findings in the Perturb-seq data, mild hypoxia during this developmental period reduced pial collateral abundance (**Figure 5l**). The scRNA-seq identified a similar panel of EC subtypes as in the Perturb-seq data, and control genes in the glycolysis pathway (*Ldha*, *Aldoa*, *Gapdh*) were upregulated as expected ^94,98^ (**Figure S7**). Mild hypoxia also produced the opposite shift in overall EC subtypes observed when hypoxia signaling genes were knocked down in the Perturb-seq experiment. Specifically, composition shifted away from arterial states with the same inflection point between pre-artery tip cells and peri-arterial capillaries. In addition, the strongest subtype expansions were the two tip cells (**Figure 5m, compare with** **Figure 5c**). Thus, the effect of lowered environmental oxygen, at both the tissue and EC subtype levels, supports the predicted role of hypoxia genes as artery repressors.

Projecting the hypoxia and normoxia datasets onto the Perturb-seq-derived program space and calculating differences in program usage between the two conditions in all ECs showed 44 programs up in hypoxia and 49 down (**Table S10**), including the expected increase in Program 18-HIF-dependent glycolysis (**Figure 5n, top row**). Program 6-Canonical WNT was also up (**Figure 5n**). Consistent with hypoxia affecting tip cell subtypes, pre-artery tip cell-enriched Programs 11 and 47 were increased in hypoxia and artery programs (Programs 65, 68, and 20) were decreased (**Figure 5n**). Thus, the same HIF- and WNT-linked programs that oppose arterial specification in the Perturb-seq screen also respond to environmental hypoxia.

Finally, we returned to the cross-species analysis and considered whether these same gene programs were differentially expressed between guinea pig and mouse, as we had observed for the artery repressors (**Figure 4a**). Indeed, when calculating programs for all guinea pig ECs combined, artery programs (Programs 65, 68, and 20) were expressed more highly, whereas Program 18-HIF-dependent glycolysis, Program 6-Canonical WNT, and tip cell Programs 11 and 16 were reduced (**Figure 5n, bottom row; Table S11**).

Together, these data support a model in which local tissue hypoxia occurring during developmental growth activates HIF- and WNT-linked gene programs that retain ECs in an angiogenic and pre-artery tip cell state and inhibit their transition toward the arterial side of the EC continuum, balancing the artery development forces present (**Figure 5j**). In guinea pig ECs, reduced sensitivity of these programs during development may lower the barrier to tip-to-artery specification and result in more artery ECs, thereby favoring collateral artery formation (**Figure 5o**).

## Discussion

Our study combined comparative single-cell genomics with an *in vivo* Perturb-seq screen and an *in vivo* collateral development assay to causally link species-divergent gene expression to collateral artery formation. We identified several repressive programs whose attenuation promotes arterial EC specification. Guinea pig ECs express lower levels of genes related to hypoxia response, TCA cycle, WNT signaling, and proteostasis relative to mice, and the Perturb-seq screen demonstrated that partial knockdown in mouse brain ECs increases artery specification at the single-cell level. At the tissue level, in vivo inhibition of four repressor pathways (*Fzd4/6*, *Idh2/Mdh2/Ogdh*, *Hspa5*, and *Creb3l2*) increased pial collateral formation by up to 50%, establishing functional relevance of these pathways in collateral development. At the mechanistic level, repressor knockdown most frequently reduced *Esm1+* pre-artery and *Apln*+ angiogenic tip cell subtypes, identifying the tip-to-artery transition as a potential barrier in collateral development. We propose that therapeutically targeting repressor pathways in ECs could be a way to increase artery EC specification and encourage collateral formation in ischemic diseases such as stroke and CAD (Figure 5o).

Our findings are consistent with previous work suggesting that native collateral abundance is genetically regulated by developmental genes in ECs. In mice, pial collateral number differs widely across strains, and fine mapping of the C57BL/6 versus BALB/cBy difference identified *Rabep2* as a natural variant that alters embryonic collaterogenesis ^31^. Broader Collaborative Cross mapping showed that collateral number varies across a much wider genetic panel and is a highly polygenic trait, with multiple contributing loci ^16^. Our study advances this framework from genetic mapping to high-throughput causal discovery of endothelial regulators contributing to the exceptional collateralization of guinea pigs.

The guinea pig collateral phenotype may have been influenced by the species’ genetic evolution at high elevations in the Andes Mountains. Genetic studies of human populations and other vertebrates from very high altitudes have identified associations with variants in genes of the hypoxia response pathway, especially *EPAS1* and *EGLN1* ^97,99,100^. Interestingly, some of these variants limit hypoxia responsiveness, presumably protecting carriers from the effects of chronic hypoxic signaling. Although guinea pigs have not been included in these genetic studies, they share physiological hallmarks of reduced hypoxia responsiveness, including high-affinity hemoglobin, moderate erythrocytosis under chronic hypoxia, and a blunted carotid body response to hypoxia ^101–104^. This raises the possibility that the lower expression of hypoxia pathway genes in guinea pig ECs reflects genetic selection during their evolution in the Andes Mountains. If so, reduced activation of hypoxia response genes during the low oxygen window of embryonic development could tip the balance away from tip cell angiogenesis toward artery development and collateral formation (Figure 5o).

Our data suggest that tip cells sit at a decision point in artery EC specification and collateral formation. Lineage tracing in the mouse and live imaging in zebrafish have demonstrated that pre-artery tip cells form arteries in the retina, heart, and intestine, with Notch*, Eph/ephrin, Dach1,* and *Itgb1* positively influencing their decision to differentiate and/or migrate into growing arteries ^62,65–67,69,105^. At first glance, our identification of HIF–VEGF-linked programs as artery repressors may seem paradoxical, because hypoxia and VEGF signaling have been shown to promote collateral formation. Indeed, Aghajanian et al. observed that chronic systemic hypoxemia over several weeks results in increased collateral flow in the adult heart and brain, increased *Hif2a, Vegfa, Rabep2* and related angiogenic genes, and that this response is inhibited by knockdown of *Vegfa*, *Flk1/Kdr*, or *Cxcr4* or loss of *Rabep2* ^32,33^. Kumar et al. further showed that developmental pial collaterals form through *Kdr*-dependent extension of pre-existing artery tips along microvascular tracks. Our data do not argue that HIF or VEGF signaling is inhibitory throughout collateral formation. Rather, they suggest that collateral growth requires stage- and EC state-specific tuning of these pathways. Transient, lower level or spatially restricted HIF–VEGF activity may be necessary to establish and guide motile tip, pre-artery, or artery-tip populations in hypoxic or ischemic tissue, whereas persistent activation may retain ECs in angiogenic/pre-artery tip states and delay their maturation into arteries.

Consistent with the interpretation that they can inhibit artery EC specification cell autonomously, WNT and HIF–VEGF genes have emerged in other systems as antagonists of artery specification. In the coronaries during heart development, WNT activity maintains ECs in a proliferative state, and *Fzd4* loss accelerates their arterialization by enabling cell cycle exit^35^. Also in the heart, HIF activation promotes angiogenic sprouting while delaying the maturation of arteries ^106^. Finally, in the retina, VEGF-activated angiogenesis at the vascular front where tip cells reside antagonizes shear stress-induced artery specification to provide a balanced vasculature ^107,108^. In that study, EC *Kdr* deletion produced ectopic arteries, leading the field to propose that angiogenesis and arteriogenesis oppose each other ^76,105,108^.

Limitations: Discoveries reported here rely primarily on transcriptional data and focus on differences between guinea pigs and mice. Future studies could incorporate proteomic data and extend the screening to additional pathways that may be involved in collateral formation in both species, e.g., genes upregulated in mice and humans after stroke or myocardial infarction. Furthermore, while four repressor pathway groups were validated at the tissue level during collateral artery development, many of the 41 identified repressors remain untested for this specific feature *in vivo*. Testing more pathways, and even combining them, could increase our understanding of their role and lead to more effective approaches for promoting collaterals. Finally, all mechanistic experiments were conducted in neonatal mouse brain endothelium under normoxic conditions. Future studies will move to the translational question of whether attenuating these repressive programs can restore collateral formation in adult tissues during injury conditions.

## Methods

### Animal models, gestational staging, and ethics

All experiments used mouse or guinea pig tissues. No human participants, clinical specimens, or field-collected samples were included. All animal procedures were performed under Institutional Animal Care and Use Committee (IACUC) approved protocols.

Mouse embryonic stages were assigned by timed mating, with the day of vaginal plug detection designated E0.5. Guinea pig gestational stages were assigned from Charles River timed-breeding records, with embryo length and gross heart size recorded at dissection as secondary morphometric support. For cross-species developmental comparisons, heart stages were aligned based on coronary artery smooth-muscle coverage and branching morphology, as detailed below and summarized in Table 1. Perturb-seq and validation experiments used neonatal mice during the first postnatal week, with AAV delivery at P0-P2 and tissue harvest at P7-P9.

### Data availability

Raw and processed sequencing data generated in this study, including embryonic mouse and guinea pig single-cell/multiome datasets, brain endothelial single-cell RNA-seq datasets, *in vivo* Perturb-seq datasets and neonatal hypoxia single-cell RNA-seq datasets, will be deposited in GEO/SRA under accession numbers to be added before publication. Spatial transcriptomic data generated by Xenium will be deposited in an appropriate public repository, with accession numbers or persistent identifiers to be added before publication. Private reviewer-access links will be provided during peer review.

### Code availability

All computational analyses were performed using custom scripts available at https://github.com/ifanirene/collateral-artery-perturbseq-paper. Interactive figures including program annotations and perturbation-effect volcano plots for all gene programs are available at the <u>manuscript resources page</u>.

### Collateral artery phenotyping

#### Whole mount immunolabeling and imaging of coronary arterial networks

Embryonic guinea pig hearts were collected at the indicated gestational stages, fixed in 4% paraformaldehyde (PFA; Electron Microscopy Sciences, 15714) at 4°C for 1 h, washed in PBS, and stored in PBS containing 0.01% sodium azide (Sigma-Aldrich, S8032) until processing. Hearts spanning GD 25-38 were collected for developmental analysis.

Whole-heart immunofluorescence was performed following a modified iDISCO+ protocol ^109,110^. Briefly, fixed hearts were dehydrated through a graded methanol/ddH₂O series (20%, 40%, 60%, 80%, 100%, 100%), each step for 1 h with agitation, then incubated in 66% dichloromethane (DCM; Sigma-Aldrich, 34856) / 33% methanol. After a methanol wash, hearts were bleached overnight at 4°C in 5% hydrogen peroxide (Sigma-Aldrich, 216763) in methanol, then rehydrated through the reverse methanol/ddH₂O series. Hearts were next permeabilized in PBS containing 0.2% Triton X-100, incubated at 37°C for 2 days in 20% DMSO / 2.3% glycine (Sigma-Aldrich, G7126) / 0.2% Triton X-100/PBS, and blocked at 37°C for 2 days in 10% DMSO / 6% normal donkey serum (Jackson ImmunoResearch, 017-000-121) / 0.2% Triton X-100/PBS. Primary antibodies were diluted in PBS containing 5% DMSO, 3% normal donkey serum, 0.2% Tween-20, and 0.1% heparin (Sigma-Aldrich, H3393) and incubated at 37°C for 4-14 days. Arterial networks were visualized with Cy3-conjugated SMA antibody (1:300; Sigma-Aldrich, C6198), together with endothelial markers as appropriate. Washes after antibody incubations were performed in PBS containing 0.2% Tween-20 and 0.1% heparin. Before optical clearing, samples were embedded in 1% low-melting agarose (Sigma-Aldrich, A9414) in PBS, dehydrated again through graded methanol, incubated in 66% DCM / 33% methanol, washed in 100% DCM, and cleared in ethyl cinnamate (ECi; Sigma-Aldrich, 112372). Cleared hearts were kept at room temperature in the dark until imaging.

Cleared hearts were imaged on a LaVision BioTec Ultramicroscope II light-sheet microscope using Imspector Pro software, with the sample in a quartz cuvette filled with ECi. Imaging used an Olympus MVX10 zoom body with a 2X objective; z-stacks spanning the full heart were acquired for 3D reconstruction. Maximum-intensity projections were generated in Fiji/ImageJ, and 3D rendering was performed in Imaris ^111^. Representative images were lightly adjusted for contrast or sharpness for display only; collateral identification and quantification were performed on the underlying image data.

#### Three-dimensional tracing and quantification of coronary collaterals

Collateral quantification was performed as described ^112^. Sequential light-sheet image stacks were imported into NIH Fiji/ImageJ and prepared for semi-automated artery tracing by converting the stacks to 8-bit images and reducing the resolution to one-fourth of the original size. Coronary arterial trees were traced using the Simple Neurite Tracer plugin, or the updated SNT plugin after discontinuation of Simple Neurite Tracer, by placing seed points along the length of SMA-positive arterial vessels. After all SMA-positive arteries in each coronary branch were accounted for, traces were isolated with the Fill Out function to generate a three-dimensional outline of the arterial structure for downstream quantification.

Coronary collateral arteries were defined as direct artery-to-artery bridges connecting two distinct coronary arterial branches. In whole mount preparations, candidate collaterals were identified as continuous SMA-positive segments bridging adjacent coronary branches rather than terminating within the capillary bed. Collateral abundance was quantified per heart by counting all identifiable artery-to-artery bridges from whole mount projections together with corresponding 3D reconstructions. Developmental tracings and quantification were performed across GD28-GD38 to define the emergence and expansion of coronary collaterals.

#### Imaging and quantification of pial collateral arteries

Whole brains were collected from mice and guinea pigs at the indicated developmental stages or experimental endpoints. Following euthanasia, the skull was carefully removed, and brains were gently lifted out with meninges and cortical surface intact. Samples were fixed in 4% PFA at 4°C, washed in 1X PBS, and processed as whole mounts. Brains from control and experimental groups were collected and processed in parallel whenever possible.

Fixed brains were permeabilized in 1X PBST and incubated overnight at 4°C with Cy3-conjugated SMA antibody (1:300; Sigma-Aldrich, C6198) to label smooth-muscle-covered arteries. After extensive PBS washes, brains were maintained in PBS or anti-fade mounting medium and imaged as whole mounts.

Brains were oriented dorsal side up and imaged on a Zeiss LSM 980 confocal microscope. Low-magnification tile scans or z-stacks captured the overall pial arterial pattern, followed by higher-magnification images of the cortical watershed region to resolve individual collateral connections. Imaging settings were matched across conditions within each experiment. Image stacks were processed in Fiji/ImageJ to generate maximum-intensity projections, with tile fields stitched when needed to reconstruct the full pial network.

Pial collateral arteries were defined as short artery-to-artery connections bridging adjacent arterial territories within the cortical watershed region, distinguished from distal branches by their position between neighboring arterial trees and their morphology within the surface arterial plexus. Collateral abundance was quantified manually from projected images per hemisphere or per brain, as specified in the relevant figure legends. The same anatomical criteria were applied across species and experimental conditions throughout the study.

### Single-cell and spatial profiling of developing vasculature

#### Cross-species developmental stage alignment and sampling design

Cross-species atlas sampling was anchored to the onset and maturation of coronary artery smooth-muscle coverage. Embryonic hearts were collected from mice at E13.5, E15.5, and E17.5 and from guinea pigs at GD25, GD32, and GD35, corresponding to stages before coronary arterialization, during arterialization and collateral initiation, and after collateral initiation, respectively. To test whether species-divergent endothelial programs were shared across organs, late-stage brain endothelial datasets from mouse E18/P0 and guinea pig GD35 were analyzed in parallel. These time points were selected to align with the heart stage showing the clearest arterial-state divergence while maintaining comparable developmental anatomy. Across all organs, stages, and species, the atlas comprised more than 100,000 total cell- or nucleus-derived profiles.

#### Single-nucleus multiome library preparation from embryonic hearts

Embryonic hearts were flash-frozen after dissection and cryostored until processing. Frozen hearts were processed for paired single-nucleus RNA and ATAC-seq (snMultiome) following a nuclei-isolation workflow adapted from ^113^. Tissue was minced on dry ice and mechanically dissociated with a pre-chilled Dounce homogenizer (Millipore Sigma, D9938) in chilled homogenization buffer containing 260 mM sucrose, 30 mM KCl, 10 mM MgCl₂, 20 mM Tricine-KOH (pH 7.8), 1 mM DTT, 0.5 mM spermidine, 0.15 mM spermine, 0.3% NP-40, 60 U/mL RiboLock RNase inhibitor (Thermo Fisher Scientific, EO0381), 1% BSA, and cOmplete Protease Inhibitor Cocktail (Roche, 04693116001). Homogenates were filtered sequentially through 100-µm and 70-µm strainers and purified by iodixanol density gradient centrifugation. Purified nuclei were resuspended in ATAC-compatible buffer (10 mM Tris-HCl pH 7.5, 10 mM NaCl, 3 mM MgCl₂, 0.1% Tween-20, 1% BSA, 1 U/µL RNase inhibitor), filtered through a 40-µm strainer, counted by Trypan Blue exclusion, and loaded onto the 10x Genomics Chromium Controller using the Chromium Single Cell Multiome ATAC + Gene Expression kit (PN-1000283). Libraries were indexed using Single Index Kit N Set A (ATAC, PN-1000212) and Dual Index Kit TT Set A (RNA, PN-1000215), quality-checked with the Agilent Bioanalyzer High Sensitivity DNA kit and Qubit 1X dsDNA HS assay, and sequenced on Illumina NextSeq/NovaSeq instruments to targets of approximately 20,000 RNA reads and 25,000 ATAC reads per nucleus.

#### Endothelial cell enrichment and single-cell RNA-seq from brains

Mouse E18 and P0 brains and guinea pig GD35 brains were processed fresh as individual biological replicates. Whole brains were micro-dissected in ice-cold PBS, minced with a razor blade, and dissociated in pre-warmed buffer containing collagenase IV (500 U/mL;

Worthington, LS004186), dispase (1.2 U/mL; Worthington, LS02100), DNase I (32 U/mL; Worthington, LS002007), and 10 mM HEPES in DMEM. Samples were incubated at 37°C for 10 min and triturated with wide-bore tips until no visible clumps remained. Cell suspensions were filtered through 40-µm strainers, rinsed with DMEM + 5% FBS, and centrifuged at 400g for 10 min at 4°C, then washed once in cold staining buffer (PBS + 2% FBS + 1 mM EDTA). Dissociated cells were stained with CD31-APC (BioLegend, clone MEC13.3, 102509; 1:100) for 20-30 min protected from light, washed, filtered through a 40-µm strainer, and stained with SYTOX Blue (Thermo Fisher Scientific, S34857; 1:1,000) immediately before sorting. FACS was performed on a Sony MA900 sorter using FSC/SSC, singlet, live (SYTOX Blue-), and CD31-APC+ gates. Sorted cells were collected into tubes containing 10 µL FBS and 5 µL RiboLock RNase inhibitor, kept on ice, and processed immediately; dissociation, staining, sorting, and 10x capture were completed within 4 h. Single-cell libraries were prepared from freshly sorted cells using the Chromium Next GEM Single Cell 3’ Kit v3.1 (10x Genomics, PN-1000128), targeting 5,000-10,000 endothelial cells per replicate, and sequenced on Illumina NovaSeq instruments to obtain >20,000 mean reads per cell.

#### Spatial gene expression profiling of embryonic hearts by Xenium

Spatial validation was performed on one heart per species collected at mouse E18 and guinea pig GD40. We designed a homology-aware Xenium panel of 299 genes targeting conserved transcript regions shared between mouse and guinea pig, including arterial identity markers, canonical pro-arterial ligands and receptors, and metabolic and stress-response genes prioritized from the cross-species single-cell analyses. Hearts were embedded, sectioned, and processed according to the standard 10x Xenium workflow ^114^.

For computational analysis, Xenium region outputs were imported into Seurat v5.3.0 with “LoadXenium” and cells were filtered to those with 100-2,000 transcripts and at least 50 detected genes. The four region outputs (two per species) were merged and sketch-integrated using CCA, then clustered and annotated by marker expression against the scRNA-seq-defined arterial continuum. Endothelial cells were identified by expression of at least two of eight canonical EC markers (*Flt1*, *Kdr*, *Tek*, *Egfl7*, *Klf2*, *Klf4*, *Dach1*, *Notch1*). Raw transcript counts were summed within each replicate unit to generate endothelial pseudobulk matrices, and differential expression between species was tested with DESeq2 v1.42.0 ^115^ using the design formula ∼ region + species.

### Cross-species bioinformatic analysis

#### Guinea pig genome reference construction and sequencing data processing

Multiome sequencing reads were aligned using Cell Ranger ARC v2.0.2. Mouse libraries were aligned against the pre-built mm10 reference (refdata-cellranger-arc-mm10-2020-A-2.0.0; GRCm38/Ensembl 98). Guinea pig libraries were aligned against a custom reference built with “cellranger-arc mkref” from the NCBI mCavPor4.1 assembly (GCF_034190915.1; annotation release RS_2024_02). The GTF annotation was pre-filtered to retain protein-coding and lncRNA biotypes only, and a JASPAR2024 Core vertebrate non-redundant motif file was supplied for transcription factor motif annotation at chromatin peaks ^116^. The resulting reference contained STAR-based indices for RNA alignment and BWA-based indices for ATAC alignment; build files are available upon request. Brain EC single-cell libraries were aligned against the mm10-2020-A reference (GENCODE vM23/Ensembl 98) using Cell Ranger v7.

#### Cross-species atlas integration and endothelial annotation

Filtered 10x gene-expression matrices were processed in Seurat v4.4.0 ^117^, retaining genes detected in at least three cells. Each sample was log-normalized, variable features were selected, and low-quality cells were excluded (heart mouse: >750 genes, >1,000 UMIs, mitochondrial fraction <10%; brain: >1,000 genes, >1,000 UMIs, mitochondrial fraction <12.5%). Doublets were scored with scDblFinder v1.17.2 ^118^ in cluster-aware mode with a 2.5% expected doublet rate and cells with score ≥ 0.2 were excluded.

Guinea pig gene symbols were remapped to mouse nomenclature using a 1:1 ortholog table from Ensembl BioMart ^119^. Mouse and guinea pig objects were then integrated with Seurat’s “FindIntegrationAnchors” or “IntegrateData” workflow, with highly variable genes identified within each sample and shared integration features selected across all six samples. The integrated heart atlas was reclustered after PCA and graph-based neighbor finding, and the endothelial compartment was isolated for focused reanalysis. In a subsequent endothelial-only analysis, the six heart samples were reintegrated using 1,000 selected integration features, and the endothelial object was reclustered after PCA (20 components), neighborhood graph construction (dimensions 1-7), clustering at resolution 0.4, and UMAP (n.neighbors = 50). Cells with scDblFinder.score ≥ 0.2 were excluded and only scDblFinder-called singlets were retained. Endothelial subtype labels, spanning large artery, arteriole, peri-arterial capillary, angiogenic tip, pre-artery tip, venous, and cycling states, were assigned based on canonical marker expression (*Gja5*, *Gja4*, *Cxcr4*, *Aplnr*, *Apln*, *Nr2f2*, *Nrp2*, *Vwf*, *Bmx*).

The brain EC comparison followed the same overall procedure: one Seurat object was created per sample, initial QC applied (>1,000 genes and >1,000 UMIs for all samples; mitochondrial fraction <12.5%), endothelial clusters isolated and reintegrated in the shared ortholog space, and cells with scDblFinder.score ≥ 0.2 excluded. A higher-stringency brain endothelial subset (nFeature_RNA > 2,000, nCount_RNA > 3,000, percent.ribo > 5) was reclustered and annotated into arterial, pre-artery, capillary, cycling, and venous endothelial states. Cell cycle phase was scored with “CellCycleScoring” using gene sets from ^120^ in both heart and brain datasets.

#### Continuous arterial identity scoring

Arterial identity was quantified as a continuous per-cell score rather than by binary classification, because arterial ECs represented a minority of total ECs and many cells occupied intermediate capillary-to-arterial states. Tissue-specific top-50 marker lists for arterial states (large artery plus arteriole) from heart and brain were scored with UCell v2.2.0 ^121^, a rank-based method that summarizes how strongly each cell expresses a given marker signature. Representative shared markers included *Gja4*, *Gja5*, *Igfbp3*, *Unc5b*, *Vegfc*, and *Efnb2*. Score distributions were compared across species within matched heart and brain endothelial subsets.

#### Differential expression and pathway analysis across species

Cross-species differential expression was performed within matched endothelial-cell comparisons across the processed heart and brain EC atlases, restricted to 1:1 ortholog pairs. For pathway-level analysis in each organ, we rebuilt working matrices from raw count layers, renormalized with Scanpy v1.11.3 ^122^ (“normalize_total” to 10,000 counts, then “log1p”), and scored the 50 MSigDB Hallmark gene sets (v2024.1) with decoupler v2.1.1 AUCell (‘tmin = 5’) ^123,124^. Pathway activity was then compared between guinea pig and mouse using Scanpy’s Wilcoxon rank-sum framework, and pathway log2 fold-changes with Benjamini-Hochberg-adjusted p-values were extracted. Heart and brain pathway tables were merged to assess directional concordance across organs; the cross-organ comparison highlighted pathways with FDR < 0.01 and |log2 fold-change| ≥ 0.25 in both organs.

#### Intercellular signaling inference with CellChat

We used CellChat v2.1.2 ^48^ to test whether ECs in the heart atlas received globally increased canonical pro-arterial signaling from surrounding cardiac cell populations. One CellChat object was created per species from the annotated heart Seurat object using the mouse interaction database, and communication strengths were inferred using tri-mean average expression with a minimum sender-receiver cell threshold of three. To compare canonical pro-arterial signaling, we summed communication strengths across all ligand-receptor pairs per pathway for each sender-to-EC comparison and assessed whether pathways such as VEGF, NOTCH, and TGFβ were elevated in guinea pig. For receptor candidate nomination, we applied “netMappingDEG” to link each EC-targeted interaction to the direction and magnitude of the relevant ligand and receptor differential expression, and ranked receptors by the strongest EC-targeted interaction per receptor after filtering for concordant ligand-receptor differential expression between species.

#### Single-cell metabolic pathway activity inference with scFEA

We used single-cell Flux Estimation Analysis (scFEA) ^52^ to ask whether transcript-level changes in metabolic genes were accompanied by coordinated differences in inferred pathway activity across cardiac cell types. scFEA estimates the relative activity of curated metabolic modules from enzyme-encoding gene expression, constraining connected reactions to remain metabolically consistent. Results were interpreted as directional indicators of relative glycolytic and TCA cycle activity across species and cell types, rather than as absolute biochemical fluxes or metabolite concentrations.

#### Chromatin accessibility processing and peak-gene coaccessibility scoring

Per-nucleus ATAC-seq fragment files from Cell Ranger ARC were imported into R v4.2.0 using Signac v1.10.0 ^125^. Nuclei were retained if they had 2,000-25,000 ATAC peak-region fragments, a nucleosome signal below 2.5, and a TSS enrichment score above 2. A standard Signac chromatin assay was built for mouse against the mm10 genome (EnsDb.Mmusculus.v79 annotations); for guinea pig, a custom BSgenome data package was constructed from the NCBI GCF_034190915.1 genomic FASTA using the UCSC faToTwoBit utility and “BSgenome::forgeBSgenomeDataPkg()”; gene annotations were imported from the NCBI GTF for the same assembly (release RS_2024_02, pre-filtered to protein-coding and lncRNA biotypes). Because guinea pig NCBI chromosome identifiers contain periods in the accession format, chromosome names were reformatted before downstream processing. This package is available upon request.

Peak-to-gene coaccessibility scores were computed using Cicero v1.3.9 ^126^ (cole-trapnell-lab/monocle3 branch). For each species, nuclei were first dimensionally reduced by latent semantic indexing in Monocle3 v1.3.1 ^127^ followed by UMAP embedding, then passed to “make_cicero_cds” and “run_cicero” with default parameters. To account for sequencing depth differences, mouse ATAC fragment counts were downsampled by binomial thinning (k = 100) to match guinea pig in-peak fragment depth before fitting the Cicero model. Peak-gene pairs with a coaccessibility score ≥ 0.54 were retained as input for CellOracle gene regulatory network construction.

#### Transcription factor perturbation scoring with CellOracle

To nominate transcription factors for the Perturb-seq library, we applied CellOracle v0.18.0 ^128^ (Python 3.8.18, scanpy v1.9.6) to guinea pig and mouse embryonic heart endothelial cells independently. A custom TSS annotation for the guinea pig genome was built from the NCBI GFF3 annotation file (release GCF_034190915.1-RS_2024_02) by computing strand-aware promoter windows (1,000 bp upstream / 100 bp downstream of each TSS per transcript), merging overlapping windows per gene with pybedtools v0.9.1 ^129^, and registering the resulting 27,948-entry BED file with CellOracle utilities. Mouse analysis used CellOracle’s built-in mm10 reference.

The base GRN for each species encoded candidate TF-target gene pairs supported by chromatin coaccessibility. CellOracle was initialized with the Cicero coaccessibility links derived above (20,699 peak-gene links for mouse; 20,659 for guinea pig), and underlying peak sequences were scanned for TF binding motifs using gimmemotifs v0.17.0 against the gimme.vertebrate.v5.0 database (false positive rate = 0.02; minimum motif score ≥ 10) ^130^, yielding a binary base GRN with 1,094 candidate TF regulators per species. Cluster-specific GRN edge weights were fitted from normalized scRNA-seq count matrices of heart endothelial cells (979 guinea pig, 1,247 mouse) spanning seven transcriptional states (Artery, Cap1-Cap4, Pre-artery, Vein). Expression values were smoothed by KNN imputation (k = 24 for guinea pig, k = 31 for mouse, each approximately 2.5% of cells) on the top 1,000 most variable genes, and GRN edge weights were estimated by bagging ridge regression across the full 3,000-gene oracle space (“oracle.get_links”, alpha = 10), fitting independent models for each cluster. Edges with p ≥ 0.001 were discarded and only the top 2,000 edges per cluster by regression coefficient magnitude were retained.

A vein-to-artery developmental trajectory was defined using diffusion pseudotime (DPT) computed in Scanpy v1.9.6 with a 30-neighbor graph and a Vein root cell, with two lineages defined: an arterial lineage (Lineage_VA: Vein → intermediate Cap clusters → Pre-artery → Artery) and a capillary lineage (Lineage_VC: Vein → Cap clusters → Cap1). Per-cell pseudotime values were smoothed onto a 40 x 40 UMAP grid by polynomial regression (degree 3) to produce a pseudotime gradient vector field; grid points with fewer than 1.8 neighboring cells (guinea pig) or 3.8 (mouse) were excluded to avoid unreliable gradient estimates in sparsely occupied UMAP regions. In silico TF knockouts were simulated for all transcriptionally active TFs (167 guinea pig, 129 mouse) by setting each TF’s expression to zero and propagating effects through the GRN for three steps (“n_propagation = 3”). Perturbation-induced cell shifts were projected into PCA space to generate a perturbation vector field, with transition probabilities estimated using 200 nearest neighbors (“sigma_corr = 0.05”). Simulations were performed across five cell-population subsets per TF (whole endothelial population, Lineage_VA, Lineage_VC, cycling capillary cells, and arterial cells). The alignment of each perturbation vector field with the pseudotime gradient field was quantified as *ps_sum*, the summed inner product across the late arterial bins (bins 6-9 of 10) within Lineage_VA. TFs with Bonferroni-adjusted p < 0.01 (one-sided paired Wilcoxon signed-rank test against permuted perturbation vector directions) were classified as predicted artery repressors. Fifty-two guinea pig TFs and 42 mouse TFs met this threshold; those with concordant significance in both species and corroborating cross-species differential expression in heart and brain ECs were prioritized for the Perturb-seq library, contributing 56 TF targets to the 162-gene screen.

### Perturb-seq library design and candidate nomination

The 162-gene perturbation library was assembled from three candidate pools derived from complementary cross-species analyses. All candidates were required to be expressed in endothelial cells and likely to act cell-autonomously, and all three pools were filtered to genes expressed at ≥10 TPM in the postnatal brain endothelial dataset.

#### Receptor candidates (50 targets)

From the EC-targeted CellChat interaction table filtered by concordant ligand and receptor differential expression (described above), receptors were ranked by their log2 fold-change between guinea pig and mouse, and the top 50 were selected after applying the ≥10 TPM filter.

#### Transcription factor candidates (56 targets)

From the statistically significant CellOracle perturbations (Bonferroni-adjusted p < 0.01, described above), the 56 TFs with the strongest predicted effect on arterial specification upon knockout in either guinea pig or mouse were selected after applying the ≥10 TPM filter. This included both predicted artery repressors (knockout promotes the arterial trajectory) and essential artery drivers (knockout deflects cells away from it).

#### Differentially expressed gene candidates (40 targets)

Candidates were ranked by concordance and magnitude of log2 fold-change between guinea pig and mouse ECs across heart and brain, prioritizing genes altered consistently in both organs. Genes in hypoxia-response, metabolic, and proteostasis/stress-response gene sets highlighted by the cross-species Hallmark pathway analysis were additionally up-weighted, as were DEGs with independent mouse genetic evidence of a vascular morphology phenotype (MP:0001614). The pool also included 16 positive and negative control genes: established regulators of endothelial arterial fate specification, flow-mediated signaling, and cell cycle ^49,75,82^. These control genes were included to benchmark screen sensitivity and calibrate the readout.

### In vivo CRISPRi Perturb-seq screen

#### Transgenic mouse lines and endothelial CRISPRi expression

CRISPRi screens were performed in compound transgenic mice carrying both *Tie2Cre* (JAX #008863) and *ROSA26-LSL-KRAB-dCas9-BFP* (JAX #026175) alleles ^58,59^, enabling endothelial-specific expression of the *KRAB-dCas9* repressor after Cre-mediated recombination. Breeding colonies were maintained on a mixed C57BL/6J background with heterozygous carriers for both alleles. Genotyping was performed by PCR on tail or ear punch DNA.

#### Guide RNA pool design

For each target gene, five gRNAs were selected from the Weissman CRISPRi mouse library (mCRISPRi-v2) ^131^, which provides ten gRNAs per gene ranked by guide quality metrics including off-target scores; the top five ranked guides were selected for each gene. For genes with alternative transcription start sites (TSS) relative to the default TSS annotations in the Weissman library, as identified by single-cell ATAC-seq data, gRNAs were designed against the relevant promoter regions using CRISPick for CRISPRi with S. pyogenes Cas9 ^132^, and the top five ranked guides per promoter were selected. The final library targeted 162 genes and comprised 889 gene-targeting gRNAs, 100 non-targeting controls from the mCRISPRi-v2 library, and 100 safe-targeting controls from the Bassik mouse library ^133^, for a total of 1,089 guides.

#### Pooled gRNA library construction and AAV packaging

The pooled gRNA expression construct was generated in an AAV transfer plasmid for packaging into a brain endothelial cell-tropic capsid, based on the AAV9-X1.1 system described by Chen et al. ^57^. The plasmid (Addgene deposit in progress) contains a human U6 promoter driving gRNA expression with Capture Sequence 1 ^60^, an EF1α promoter driving GFP, and AAV2 inverted terminal repeats. The pooled gRNA library was synthesized as a Twist oligo pool with the sequence format (GGCTTTATATATCTTGTGGAAAGGACGAAACACC + spacer (starting with G) + GTTTAAGAGCTATGCTGGAAACAGCATAGCA) and cloned into the SapI-digested backbone by Gibson assembly. Following bead purification, the assembly reaction was electroporated into ElectroMax Stbl4 competent cells (Invitrogen, 11635018), expanded overnight, and purified by maxiprep (Zymo Research D4203). Library quality was assessed by PCR amplification of the gRNA region followed by next-generation sequencing on a MiSeq instrument, confirming 100% guide representation and a library skew ratio of 2.5. AAV production was performed by the HHMI AAV Core using the corresponding brain endothelial cell-tropic packaging plasmid.

#### Retro-orbital AAV injection and brain tissue collection

Neonatal pups (P0-P1) from *KRAB-dCas9; Tie2Cre* crosses were briefly anesthetized on ice for 2-3 min and injected retro-orbitally with 10-20 µL of AAV (2 x 10¹¹ GC per pup) using a 33-gauge Hamilton syringe under a dissecting microscope. After injection, pups were warmed on a heating pad and returned to the dam. Pups were monitored daily for normal weight gain and health. Brains were harvested at P7-P9 by decapitation, and cortices were microdissected in ice-cold PBS.

#### Immunomagnetic enrichment and single-cell library preparation

Microdissected brains from 5-8 pups per replicate were dissociated using the buffer and protocol described above for brain endothelial cell preparation. After washing, cells were resuspended in cold staining buffer and incubated with EasySep Mouse FcR Blocker (STEMCELL Technologies, 18731; 40 µL/mL) followed by CD31-APC (1:100) for 20-30 min protected from light. CD31+ cells were enriched using the EasySep Release Mouse APC Positive Selection Kit (STEMCELL Technologies, 100-0033) without columns; the enriched fraction was washed, filtered through a 40-µm strainer, and stained with SYTOX Blue immediately before sorting. FACS was performed on a Sony MA900 sorter using FSC/SSC, singlet, live (SYTOX−), CD31-APC+, and GFP+ gates. Sorted cells were collected into tubes containing FBS and RiboLock RNase inhibitor, kept on ice, and processed immediately. The complete workflow, from dissociation through enrichment, sorting, and 10x capture, was completed within 4 h.

Single-cell libraries were prepared using the Chromium GEM-X Single Cell 3’ v4 workflow with direct guide capture using capture sequence 1 (10x Genomics, 4 rxns, PN-1000686; Feature Barcode Kit v4, PN-1000702). Target cell recovery was 10,000-20,000 cells per replicate. Gene-expression and CRISPR guide-capture libraries were sequenced on Illumina NovaSeq X 25B flow cells using a 28/10/10/90 cycle configuration, pooled at a 10:1 molar ratio, targeting >20,000 reads per cell for gene expression and >10,000 reads per cell for guide capture.

#### Sequencing data processing and gRNA assignment

Raw sequencing data were processed with Cell Ranger v7.1.0. Because gene-expression libraries were prepared with the 3’ v4 chemistry, the default 3’ v3 barcode whitelist was replaced with the manufacturer-supplied 3’ v4 whitelist. Gene Expression and CRISPR Guide Capture FASTQ files were processed separately and aligned to mm10-2020-A using a custom feature reference describing all gRNA oligonucleotides and direct-capture sequence 1 guides. For resequenced libraries, the deeper gene-expression matrices were merged with guide counts from the original runs, retaining barcodes present in both assays.

Filtered matrices were imported into Seurat v4, with RNA as the primary assay and guide counts as a secondary assay. Cells with 2,000-7,500 detected genes, 3,000-30,000 UMIs, and mitochondrial fraction < 10% were retained. Following log-normalization, variable feature selection, PCA, UMAP, and graph-based clustering, doublets were identified with scDblFinder v1.14.0 in batch-aware mode and excluded. Multiplicity of infection (MOI) was defined as the number of guides with > 5 UMIs per cell. For downstream Perturb-seq analyses, we retained singlet cells with at least one high-confidence guide assignment (≥ 10 guide UMIs) and MOI < 15. Clusters were annotated as endothelial and perivascular populations based on canonical marker genes.

### Screen quality control and artery-state analysis

#### Screen benchmarking and perturbation quality control

Across 10 mice, approximately 260,000 GFP+CD31+ ECs were recovered after initial sequencing and quality filtering. Screen quality was assessed at the levels of cell recovery, guide recovery, guide multiplicity, on-target repression, and calibration of perturbation–phenotype tests, consistent with quality control principles used in direct-capture and large-scale Perturb-seq studies ^37,60^. Barcode-rank plots showed a clear knee separating cell-associated from background barcodes, and library complexity indicated robust transcript capture. Approximately half of cells received a high-confidence guide assignment, with guide multiplicity centered in the intended high-MOI range (approximately 5-15 guides per cell) and robust guide UMI detection.

On-target knockdown was quantified with SCEPTRE v0.1.0 ^74^ in high-MOI mode using the gene-by-cell UMI count matrix as the response matrix, the guide-by-cell UMI count matrix as the perturbation matrix, and a curated gRNA-to-target annotation table covering 884 targeting guides across 162 genes together with 200 non-targeting guides. More than 90% of targeted genes showed significant mRNA reduction (mean knockdown efficiency approximately 40%), and batch-specific knockdown estimates were concordant (R² = 0.754). Positive-control pairs were defined as cis guide-target pairs from “construct_positive_control_pairs”, discovery testing used the trans pair set from “construct_trans_pairs”, and model calibration was assessed with “run_calibration_check” using 200,000 negative-control pairs per analysis. On-target guide-target pairs produced substantially stronger differential-expression signal than non-targeting guides, and QQ plots showed appropriate null calibration, confirming that discovery signal was not driven by model miscalibration.

As a biological benchmark, we examined *Notch1*, a canonical inducer of arterial fate ^75,82^. Averaged transcriptional changes in *Notch1* guide-assigned ECs were compared with a published bulk RNA-seq dataset showing the expected reduction in arterial markers in E14.5 coronary ECs after inducible endothelial *Dll4* conditional knockout (*Dll4* iEC-CKO; *Pdgfb*-icreERT2; *Dll4*^f/f^, tamoxifen at E13.5, tissue collected 24 h later) ^76^. DLL4 is the primary endothelial ligand signaling through NOTCH1 in pre-artery ECs, and the core transcriptional consequences of DLL4-NOTCH1 loss are conserved across coronary, retinal, and brain vasculature ^75,76^; no equivalent NOTCH1 EC-specific loss-of-function transcriptomic dataset from brain endothelium is currently available.

#### Arterial identity scoring and SCEPTRE perturbation testing

To identify perturbations that altered arterial specification, we used continuous per-cell large artery and arteriole scores as the primary screening readouts. Marker genes for each arterial state were identified with Seurat “FindAllMarkers” (test.use = “roc”, min.pct = 0.1, logfc.threshold = 0), ranked by AUC, log2 fold-change, and detection fraction; mitochondrial and ribosomal genes were excluded. The top 50 markers for each arterial state were scored in all retained endothelial cells with UCell v2.2.0, yielding two rank-based continuous artery-state scores per cell.

These scores were used as the response matrix for SCEPTRE v0.1.0 testing in high-MOI mode, retaining only barcodes shared across the guide matrix, singlet metadata, and score matrix. Extra covariates for guide assignment included mitochondrial fraction, batch, and “cell_type”; mixture-model assignment used the formula “log(grna_n_nonzero) + log(grna_n_umis) + batch + cell_type”. Perturbation effects on each artery score were then tested with two-sided conditional resampling using the formula “log(response_n_nonzero) + log(response_n_umis) + log(grna_n_nonzero) + log(grna_n_umis) + batch”, followed by “run_qc”, “run_calibration_check”, and “run_discovery_analysis”, with Benjamini-Hochberg FDR correction. Positive controls used cis guide-target pairs from “construct_positive_control_pairs”; discovery testing used trans pairs from “construct_trans_pairs”; calibration was assessed against 200,000 negative-control pairs. This continuous-score analysis constituted the primary screen for ranking perturbations as artery repressors or artery inducers.

#### Endothelial subtype composition analysis

As an orthogonal readout, cells annotated as large artery or arteriole were collapsed into a single total-artery category, and the fraction of total-artery cells among guide-assigned ECs was compared with non-targeting controls to confirm the direction of effects observed in the continuous score analysis. Independent guides targeting the same gene were used as internal consistency checks.

To understand which non-arterial endothelial states changed when artery repressors were perturbed, the score-based framework was extended to all major endothelial states. Using cell type annotations, additional marker sets were derived for pre-artery tip, vein, capillary, and angiogenic tip populations with the same marker-discovery procedure, with signature sizes adjusted for smaller capillary populations. Per-cell UCell scores for these additional states were then tested with SCEPTRE using the same singlet-cell dataset and technical covariates.

### Endothelial gene program discovery and perturbation analysis

#### Consensus NMF decomposition of the Perturb-seq expression matrix

To resolve the downstream transcriptional modules through which artery repressors altered endothelial state, we decomposed the QC-filtered Perturb-seq endothelial expression matrix into co-regulated gene programs using cNMF v1.7.0 ^85^ on raw UMI counts exported via Scanpy v1.10.1 and anndata v0.10.7. cNMF was run on the raw-count matrix without batch correction using 10,000 overdispersed genes and 200 factorization replicates; we explored a range of candidate ranks before retaining a 100-program solution for downstream analyses. Consensus clustering used a local-density threshold of 0.2 to remove unstable spectra; 4,985 of 20,000 spectra (25%) were excluded at this step, after which the clustering diagnostic showed clear diagonal structure consistent with stable program recovery. Retained outputs included the per-cell usage matrix, consensus spectra, and gene-loading tables.

#### Gene program annotation

Gene programs were annotated using ProgExplorer, an evidence-driven workflow built for this study (https://github.com/ifanirene/ProgExplorer). For each program, the top 300 loading genes defined the broader program membership, and distinctive genes were additionally identified using a uniqueness score that up-weighted genes concentrated in a single program rather than broadly shared across programs. For enrichment, the top 100 genes were queried against STRING v12.0 ^134^ (*Mus musculus*, NCBI taxon ID: 10090) for GO Biological Process terms ^135^ and KEGG pathways ^136^; terms with FDR < 0.05 and fewer than 500 background genes were retained. Structured annotation prompts incorporating high-loading genes, distinctive genes, enrichment terms, endothelial subtype context, and perturbation-response information were processed by a large language model (LLM; Anthropic Claude Sonnet 4.5) to generate concise program names and summaries. To support interpretation, program activity was also evaluated along the vein-to-artery continuum ordered by artery probabilities computed with CellRank v2.0.7 ^137^ on the Perturb-seq endothelial UMAP; this allowed discrimination of programs active at distinct stages of arterial maturation from those enriched in venous or capillary states. Final annotations were manually reviewed against gene loadings, enrichment patterns, endothelial subtype associations, and perturbation-response profiles; expert review of a random 20% sample confirmed high concordance with LLM annotations (Cohen’s κ = 0.87).

#### Program-level perturbation testing with SCEPTRE

To test whether CRISPRi perturbations altered cNMF program usage, each cell’s cNMF usage vector was normalized to sum to one, multiplied by the cell’s total gene-expression UMI count, and rounded to generate count-like program responses. These program-response matrices were tested with SCEPTRE v0.10.0 in the same high-MOI framework used for the endothelial-state analyses, using log(response_n_nonzero), log(response_n_umis), log(grna_n_nonzero), log(grna_n_umis), and batch as covariates with Benjamini-Hochberg correction at FDR 0.1. In the all-endothelial-cell analysis, 195,363 cells entered the model, 190,300 remained after cellwise QC, and all 16,200 target-program discovery pairs passed pairwise QC with minimum support thresholds of seven nonzero treated cells and seven nonzero control cells. Calibration was assessed as a required QC step by reviewing guide-assignment plots, cellwise and pairwise QC summaries, null QQ plots, and null log-fold-change distributions; median null p-values were near 0.5 globally (0.488) and within endothelial subsets (0.472-0.496), supporting acceptable calibration. Programs highlighted in figures were selected on concordance between perturbation effects and the independent biological evidence used for annotation, rather than significance alone. The same framework was also applied within endothelial subtypes containing at least 200 cells.

#### Enrichment of gene programs across endothelial subtypes

To quantify endothelial-state enrichment for each program, the cNMF usage matrix was row-normalized so that each cell’s usage vector summed to one, and the normalized matrix was loaded into a Seurat object in which each cNMF program served as a feature and each cell’s annotation served as its identity. ROC-based differential testing with Seurat FindAllMarkers (test.use = “roc”, only.pos = FALSE, min.pct = 0.1, logfc.threshold = 0.25, max.cells.per.ident = 10000) compared the usage of each program in cells of one subtype against all remaining endothelial cells, yielding bidirectional results for all 100 programs across all subtypes. Within each subtype, results were partitioned into enriched programs (average log₂ fold change > 0) and depleted programs (average log₂ fold change < 0), and each partition was ranked by a distinct priority order: enriched programs were sorted by descending AUC, descending log₂ fold change, and descending detection fraction, whereas depleted programs were sorted by ascending AUC and ascending log₂ fold change—placing the most strongly and specifically depleted programs first—with ties broken by descending detection fraction. The analysis produced a curated list of up to ten enriched and ten depleted programs for each endothelial cell state.

### Functional validation of artery repressors in neonatal brain

#### Pathway-focused CRISPRi perturbations via AAV delivery

Validation cohorts targeted the top artery-repressor classes identified in the screen: WNT receptors (*Fzd4* + *Fzd6*), TCA-cycle enzymes (*Idh2* + *Mdh2* + *Ogdh*), proteostasis/ER-stress factors (*Hspa5*, *Hsp90ab1*, and *Creb3l2*). Small guide pools targeting one pathway node or module at a time were cloned into the AAV9-X1.1 vector backbone and packaged as separate validation viruses using the same production approach described above. Validation cohorts were dosed at P2 and compared with non-targeting control AAV delivered to littermates in parallel; brains were collected at P9.

#### Pial collateral artery quantification

Brains were processed for whole mount SMA immunostaining and confocal imaging as described above. Pial collateral arteries were counted manually on blinded image sets using the same artery-to-artery connection criteria defined in the phenotyping section. Statistical comparisons between each perturbation cohort and non-targeting controls were performed with two-sided Mann-Whitney U tests.

### Environmental hypoxia and collateral development

#### Neonatal normobaric hypoxia exposure

Neonatal hypoxia experiments were performed by the Perez laboratory under IACUC protocol APLAC 26889. Neonatal mice were exposed to either normoxia (21% O₂) or mild hypoxia (11% O₂) for 7 days beginning at P1. Litters were housed with the lactating dam in their home cage within a normobaric reduced-oxygen chamber in which ambient oxygen was lowered by supplemental nitrogen. Chamber O₂, CO₂, temperature, and nitrogen tank pressure were monitored routinely. Pups were assessed daily for normal breathing, pink skin color, presence of a milk spot, and righting reflex, and dams were monitored for nursing behavior and general health. At the end of the 7-day exposure, pups were removed from the chamber and processed for pial vascular imaging or brain EC isolation for scRNA-seq.

#### Pial collateral artery quantification under hypoxia

After normoxia or hypoxia exposure, brains were collected and processed for whole mount SMA immunostaining and confocal imaging of the pial arterial network as described above. Pial collaterals were counted manually per hemisphere on blinded projections using the same artery-to-artery connection criteria defined in the phenotyping section. Statistical comparisons were performed with a two-sided Mann-Whitney U test.

#### Single-cell profiling and gene program analysis under hypoxia

Brain ECs were isolated from normoxia- and hypoxia-exposed pups by the dissociation, enrichment, and FACS protocol described above for brain endothelial cell preparation, and single-cell libraries were prepared using the 10x 3’ protocol and processed with Cell Ranger v7.1.0. Seurat objects from normoxic and two hypoxic batches were log-normalized, scaled, and integrated using SCT-based anchors ^138^; the integrated object was embedded by PCA and UMAP, clustered, and annotated using canonical endothelial markers. Non-endothelial clusters were removed. EC subtype composition was compared between conditions by calculating the fraction of cells per annotated state in each condition; the proportion difference (hypoxia minus normoxia) identified shifts in endothelial subtype abundance. Gene program activity under hypoxia was quantified by projecting these cells onto the Perturb-seq-derived program space using STARcat ^139^, as described below.

### Gene program projection onto independent datasets with STARcat

To determine how the Perturb-seq-derived endothelial gene programs varied in independent datasets, specifically the neonatal hypoxia scRNA-seq and the cross-species brain EC atlas, we projected each dataset onto the fixed Perturb-seq program space using STARcat v1.0.9 ^139^. The projection reference comprised the consensus spectra matrix from the Perturb-seq cNMF run (K = 100), a 100-program × 10,000-gene matrix of transcripts-per-10,000 (TP10K)-scaled gene loadings. For each query dataset, raw UMI counts were column-standardized by dividing gene-wise by the across-cell standard deviation, and per-cell program usage scores were estimated by non-negative least squares (NNLS) with the program spectra held fixed, using scikit-learn v1.0.2’s “non_negative_factorization” with the Frobenius loss and multiplicative-update solver. The resulting usage matrices were row-normalized so that each cell’s program usages summed to one, enabling direct comparison of relative program activity across cells and conditions.

For the hypoxia dataset (25,482 cells), 9,755 of the 10,000 reference genes were present in the query. For the cross-species brain atlas (14,507 cells from mouse E18/P0 and guinea pig GD35), 6,704 reference genes were present in the shared ortholog space. In both applications, differential program usage was tested per program using two-sided Mann-Whitney U tests, with log2 fold-change, prevalence (fraction of cells with usage > 0.01), and Benjamini-Hochberg-corrected FDR computed across all 100 programs. Programs were considered significantly different in usage when FDR < 0.01 and at least 10% of cells in either condition exceeded the 0.01 usage threshold.

### Statistics and reproducibility

Statistical analyses were performed as described in the relevant Methods subsections and figure legends. Unless otherwise stated, tests were two-sided. For animal-level experiments, biological replicates were individual animals or brain hemispheres as indicated in the figure legends; for sequencing-based analyses, replicate definitions are specified in each analysis subsection. Error bars indicate mean ± s.e.m. unless otherwise stated. Exact sample sizes, replicate definitions, and p-values are reported in the figure legends and/or source data. For high-dimensional analyses, multiple testing was controlled using the Benjamini–Hochberg false discovery rate correction unless otherwise specified. Investigators were blinded to experimental condition during manual collateral quantification. No statistical method was used to predetermine sample size, but sample sizes were chosen based on prior experiments and feasibility of neonatal animal and single-cell sequencing workflows.

## Author contributions

X.F., R.Z., J.M.E. and K.R.-H. conceived the study and designed the overall experimental and analytical strategy, with input from T.Q. X.F. and R.Z. designed and performed the in vivo Perturb-seq experiments, validation experiments and single-cell analyses. X.F. led the cross-species and Perturb-seq data analyses. R.Z. led gRNA library design, cloning, screening optimization and sequencing library preparation. B.C.R., P.E.R.C., E.T., J.B. and J.A. contributed to guinea-pig developmental phenotyping, tissue processing, imaging and analysis. E.C. and M.S.C. contributed to animal experiments, tissue processing and single-cell experimental workflows. S. Alimukhamedov and A.I. performed and supervised spatial transcriptomics experiments. S. Agarwal and V.A.d.J.P. provided hypoxia experimental support and supervision. X.C., T.F.S. and V.G. contributed AAV capsids, viral production and AAV strategy. C.Y.P. and T.Q. contributed conceptual and experimental guidance. J.M.E. and K.R.-H. supervised the study. X.F., R.Z., J.M.E. and K.R.-H. interpreted the results and wrote the manuscript with input from all authors. All authors reviewed and approved the manuscript.

## Acknowledgements

We thank members of the Red-Horse and Engreitz laboratories for helpful discussions and feedback. We thank Zhe Qu (Beckman Institute CLOVER Center), Hyun Ah Yi (HHMI Janelia Viral Core) and Andrew D. Steele (Cal Poly Pomona Armamentarium Vector Core, ArmVC) for AAV vector production and support. We thank the Stanford animal care and flow cytometry core facilities for technical support.

## Materials & correspondence

Correspondence and requests for materials should be addressed to Jesse M. Engreitz or Kristy Red-Horse. Plasmids, guide libraries and other unique materials generated in this study are available from the corresponding authors upon reasonable request and completion of any required material transfer agreements.

---

**Figure S1.**
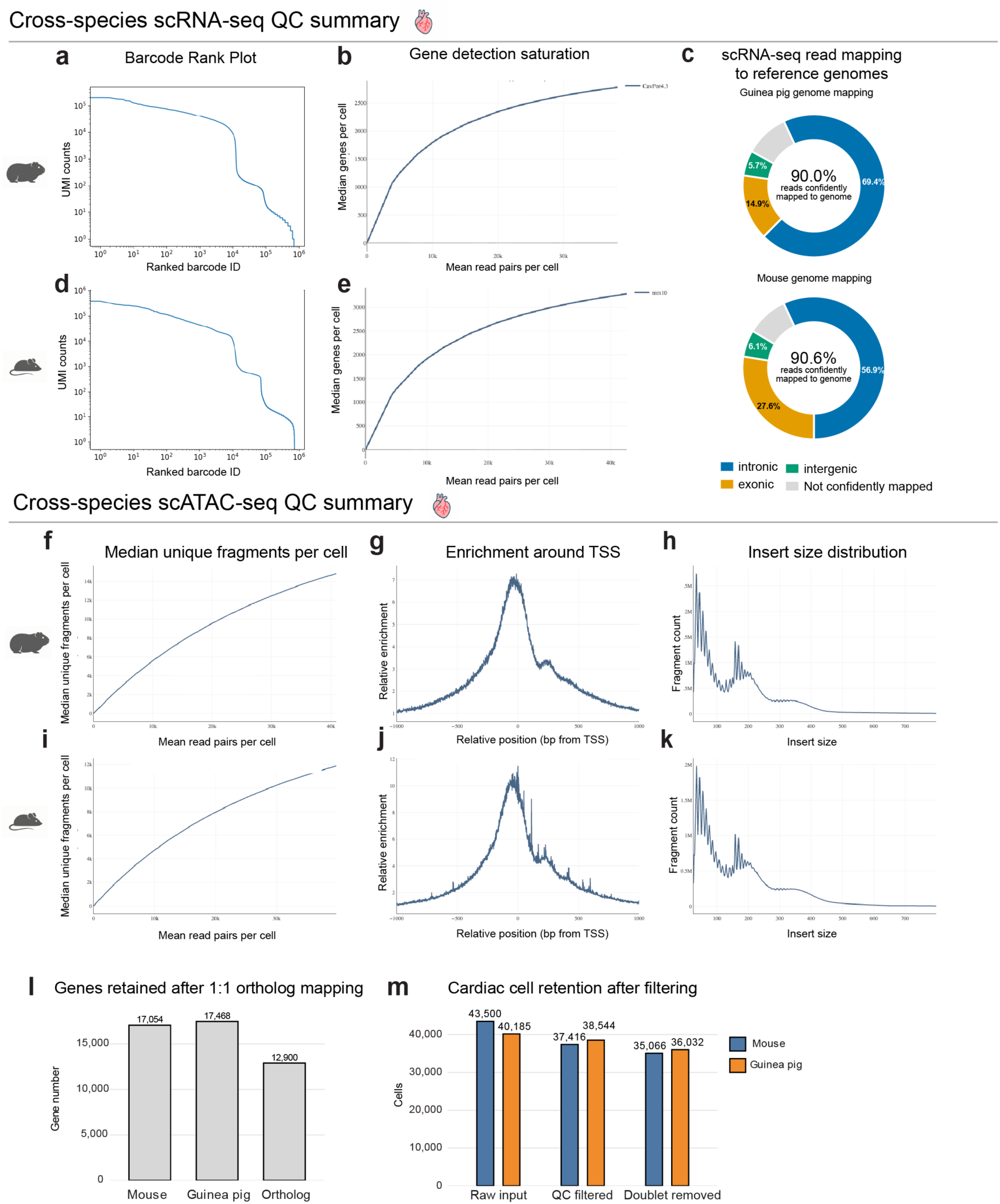
Cross-species scRNA-seq and scATAC-seq quality control summary. Across panels a-k, guinea pig data are shown in the upper row of each section and mouse data in the lower row. **a,d,** scRNA-seq barcode-rank plots show clear knees separating cell-associated barcodes from background droplets. **b,e,** scRNA-seq library-complexity curves, plotted as median genes detected per cell versus mean read pairs per cell, rise smoothly without early saturation, indicating adequate sequencing depth. **c,** scRNA-seq read-mapping summaries to the guinea pig and mouse reference genomes. Donut charts show the fractions of reads assigned to intronic, exonic, intergenic, and not-confidently-mapped categories; in both species, approximately 90% of reads mapped confidently to the genome. **f,i,** scATAC-seq library-complexity curves show increasing median unique fragments per cell with sequencing depth. **g,j,** TSS-enrichment profiles show sharp enrichment centered on transcription start sites, consistent with high-quality chromatin libraries. **h,k,** Insert-size distributions show the expected nucleosomal periodicity for ATAC-seq libraries. **l,** Genes retained after 1:1 ortholog mapping, showing the shared gene set used for cross-species analyses. **m,** Cardiac cell retention across preprocessing steps, from raw input through QC filtering and doublet removal, showing similar cell recovery between mouse and guinea pig. Together, these QC metrics support adequate library complexity, mapping quality, chromatin accessibility signal, and cell recovery across species.

**Figure S2.**
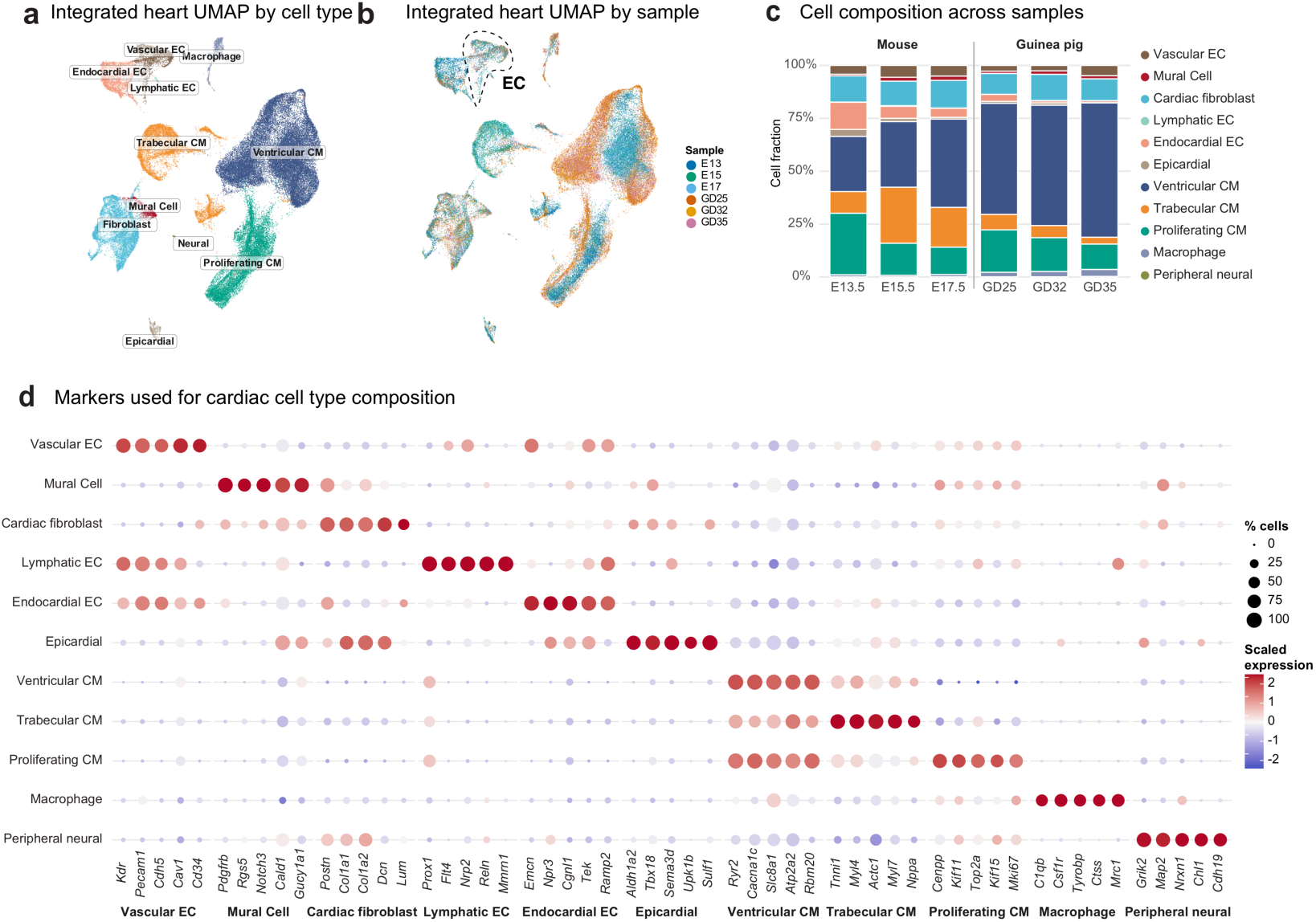
Cross-species integration of developing heart scRNA-seq and major cardiac cell type annotation. **a,** Integrated UMAP of single-cell transcriptomes from mouse and guinea pig embryonic hearts, colored by annotated major cardiac cell type. Shared clusters correspond to vascular ECs, mural cells, cardiac fibroblasts, lymphatic ECs, endocardial ECs, epicardial cells, ventricular cardiomyocytes, trabecular cardiomyocytes, proliferating cardiomyocytes, macrophages, and peripheral neural cells. **b,** Same integrated UMAP colored by developmental sample, showing that cells from different stages and species co-embed within shared cell-type clusters after integration. **c,** Major cardiac cell type composition across samples. Stacked bars show the fraction of cells assigned to each annotated population in mouse and guinea pig hearts at the indicated developmental stages. **d,** Dot plot of marker genes used for cardiac cell type annotation. Dot size indicates the fraction of cells expressing each gene, and color indicates scaled average expression within each cell type.

**Figure S3.**
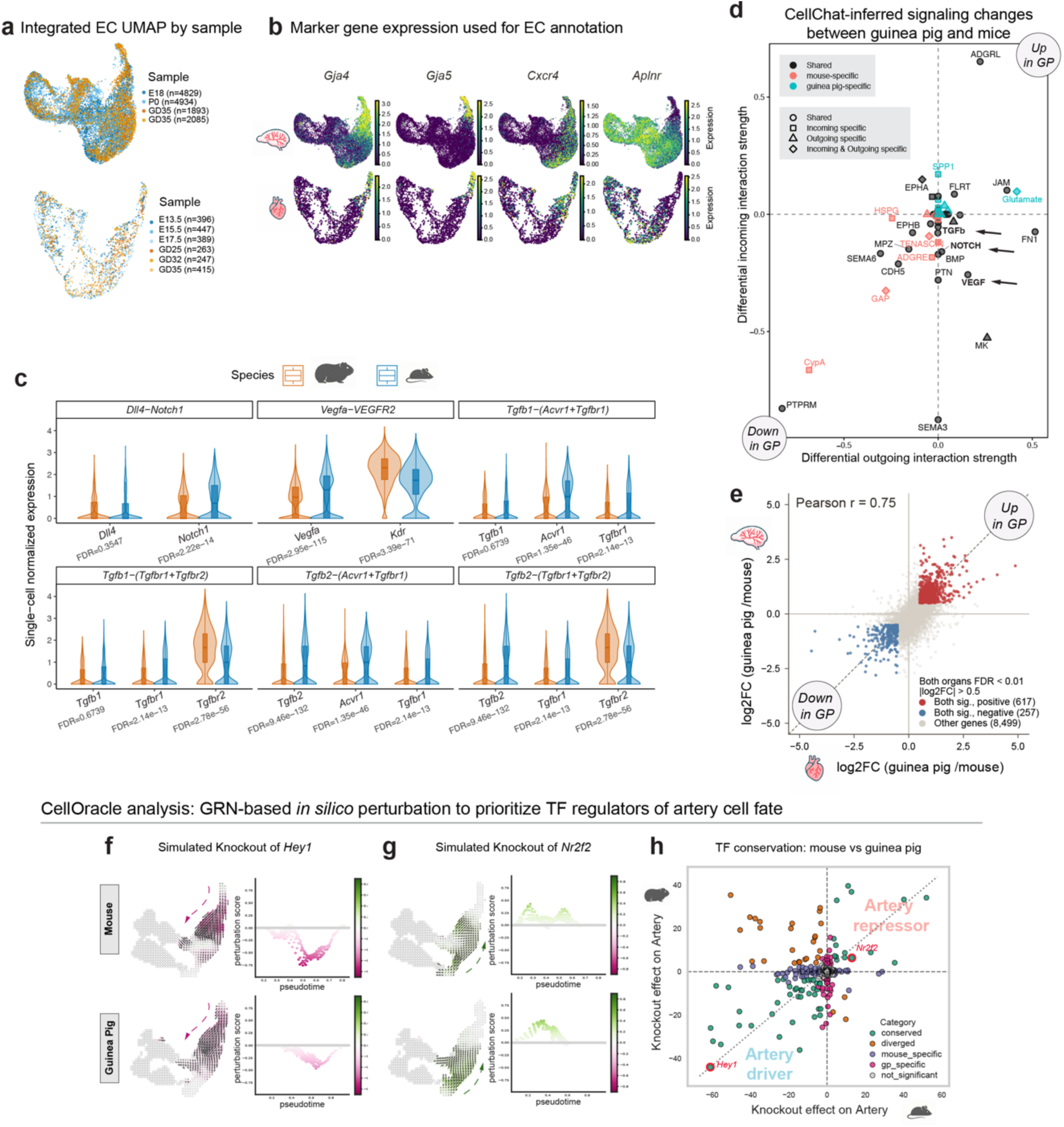
Cross-species endothelial analyses of artery identity, signaling, and gene-regulatory-network (GRN)-based transcription factor prioritization. **a**, Integrated endothelial cell (EC) UMAPs from the brain (top) and heart (bottom) datasets, colored by sample, showing that mouse and guinea pig ECs occupy the same broad EC continuum after integration. **b**, Feature plots of arterial markers used for EC annotation in the integrated brain (top) and heart (bottom) datasets. *Gja4* and *Gja5* mark arterial EC identity, whereas *Cxcr4* is enriched in pre-arterial/distal arteriole ECs and helps distinguish arterial subpopulations. **c,** Violin plots of single-cell normalized expression for genes underlying representative EC-incoming ligand-receptor interactions identified by CellChat, including DLL4-NOTCH1, VEGFA-VEGFR2, and TGFβ signaling pairs, compared between guinea pig and mouse. **d**, CellChat summary plot showing inferred signaling changes between guinea pig and mouse across major cardiac cell classes. Each point represents one signaling pathway, plotted by differential outgoing interaction strength (x axis) and differential incoming interaction strength (y axis). Pathways in the upper-right quadrant are increased in guinea pig, whereas those in the lower-left quadrant are decreased in guinea pig. **e**, Concordance of guinea pig versus mouse differential expression in ECs from heart (x axis) and brain (y axis). Each point represents one gene. The positive correlation indicates that species-biased EC transcriptional differences are strongly concordant between heart and brain. **f**, CellOracle *in silico* perturbation of *Hey1* predicts reduced arterial identity along pseudotime in both mouse and guinea pig, consistent with *Hey1* acting as an artery driver. **g**, CellOracle in silico perturbation of *Nr2f2* (*COUP-TFII*) predicts increased arterial identity, particularly along the venous branch, consistent with *Nr2f2 acting* as an artery repressor. **h**, Cross-species comparison of CellOracle transcription factor perturbation scores. Knockout effects were calculated as the signed area of Δ artery probability along pseudotime. Each point denotes one transcription factor; colors indicate conserved, divergent, mouse-specific, guinea pig-specific, or non-significant effects, separating predicted artery drivers from artery repressors.

**Figure S4.**
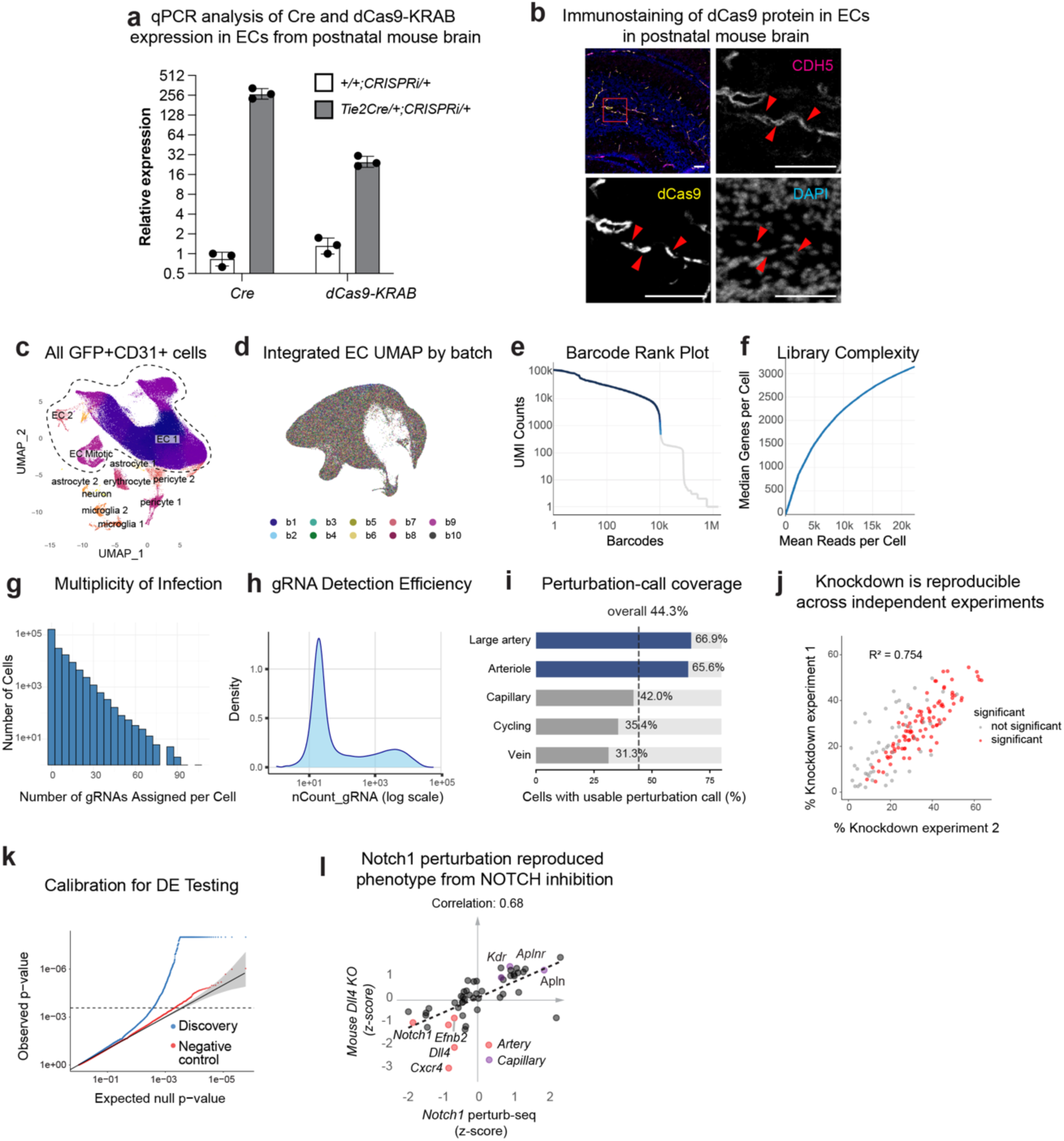
Validation and quality control of the *in vivo* Perturb-seq screen. **a**, qPCR analysis of *Cre* and *dCas9-KRAB* expression in endothelial cells isolated from postnatal mouse brains, confirming Cre-dependent *dCas9-KRAB* transgene expression in *Tie2Cre;CRISPRi* mice. **b**, Immunostaining of postnatal mouse brain vasculature showing dCas9 protein in CDH5⁺ endothelial cells. Red arrowheads indicate representative dCas9⁺ endothelial nuclei; DAPI marks nuclei. Scale bars, 50 μm. **c**, UMAP of all sorted GFP⁺/CD31⁺ cells, showing that the recovered population is predominantly endothelial, with minor non-endothelial contaminants. **d**, Integrated endothelial-cell UMAP colored by batch (10x lanes b1-b10; lanes b1, b2 were from one experiment and lanes b3-b10 were from an independent experiment using the same setup), showing good mixing across lanes and experiments. **e**, Barcode-rank (knee) plot of transcriptome UMI counts per barcode, showing a clear inflection used to define high-quality cells. **f**, Library complexity, plotted as median genes detected per cell versus mean reads per cell, indicates good complexity without early saturation. **g**, Distribution of gRNAs assigned per cell, summarizing guide multiplicity after pooled AAV delivery. **h**, Distribution of gRNA UMI counts per cell, demonstrating robust guide detection. **i**, Perturbation-call coverage across EC subtypes, showing the fraction of cells with usable perturbation calls overall and within each subtype. **j**, Per-gene knockdown efficiency is concordant between two independent experiments, shown as a correlation of per-gene knockdown estimates derived from each experiment. **k**, Calibration of differential expression testing. Negative-control tests follow the expected null distribution, whereas discovery tests show the expected enrichment of small p-values. **l,** Scatter plot of per-gene transcriptional fold changes from *Notch1* knockdown ECs from Perturb-seq (this study; x axis) versus *Dll4* iEC-CKO coronary ECs (E14.5; y axis) ^76^. Pearson r = 0.68. Highlighted genes include arterial markers down-regulated in both datasets (*Dll4*, *Efnb2*, *Cxcr4*; red) and capillary-enriched genes up-regulated in both (*Aplnr, Apln, Kdr;* purple). Concordant directionality confirms that *Notch1* knockdown in Perturb-seq from this study recapitulates the expected transcriptional hallmarks of *Notch* loss of function on arterial EC specification.

**Figure S5.**
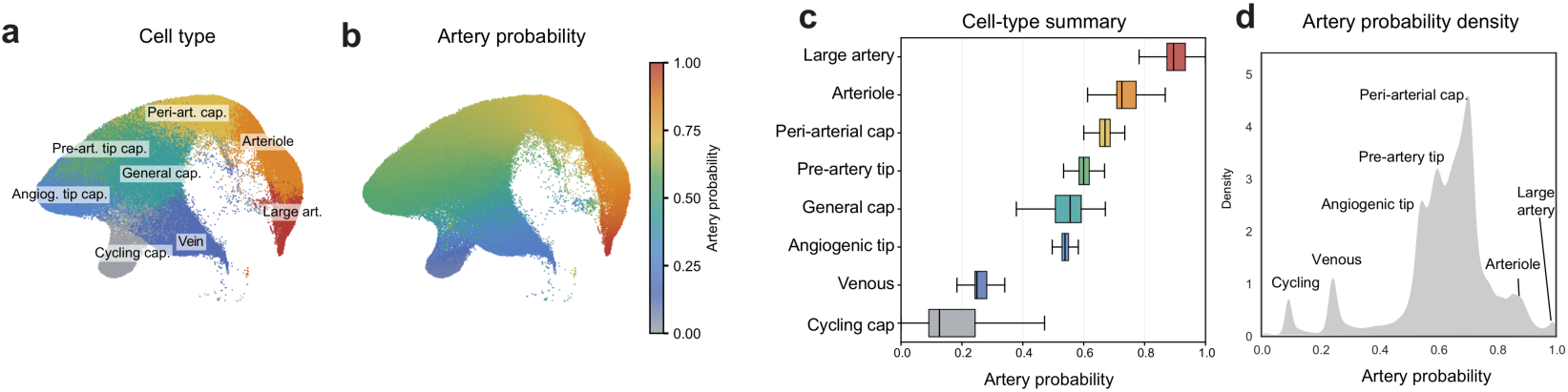
Endothelial-state ordering along the vein-to-artery continuum. Endothelial cells were ordered by artery probability to define a continuous progression from cycling/venous and capillary states to arterial states. **a,** UMAP colored by annotated endothelial state, including cycling capillary, venous, angiogenic tip capillary, general capillary, pre-artery tip capillary, peri-arterial capillary, arteriole, and large artery populations. **b,** Same UMAP colored by artery probability, showing a continuous increase from venous/capillary and tip-capillary states toward peri-arterial capillary, arteriole, and large artery identities. **c,** Horizontal boxplots summarizing artery probability for each endothelial state. **d,** Density plot of artery probability across the endothelial continuum. Together, these analyses place tip-capillary states at intermediate artery probabilities and define the trajectory framework used to interpret perturbation effects on artery specification.

**Figure S6.**
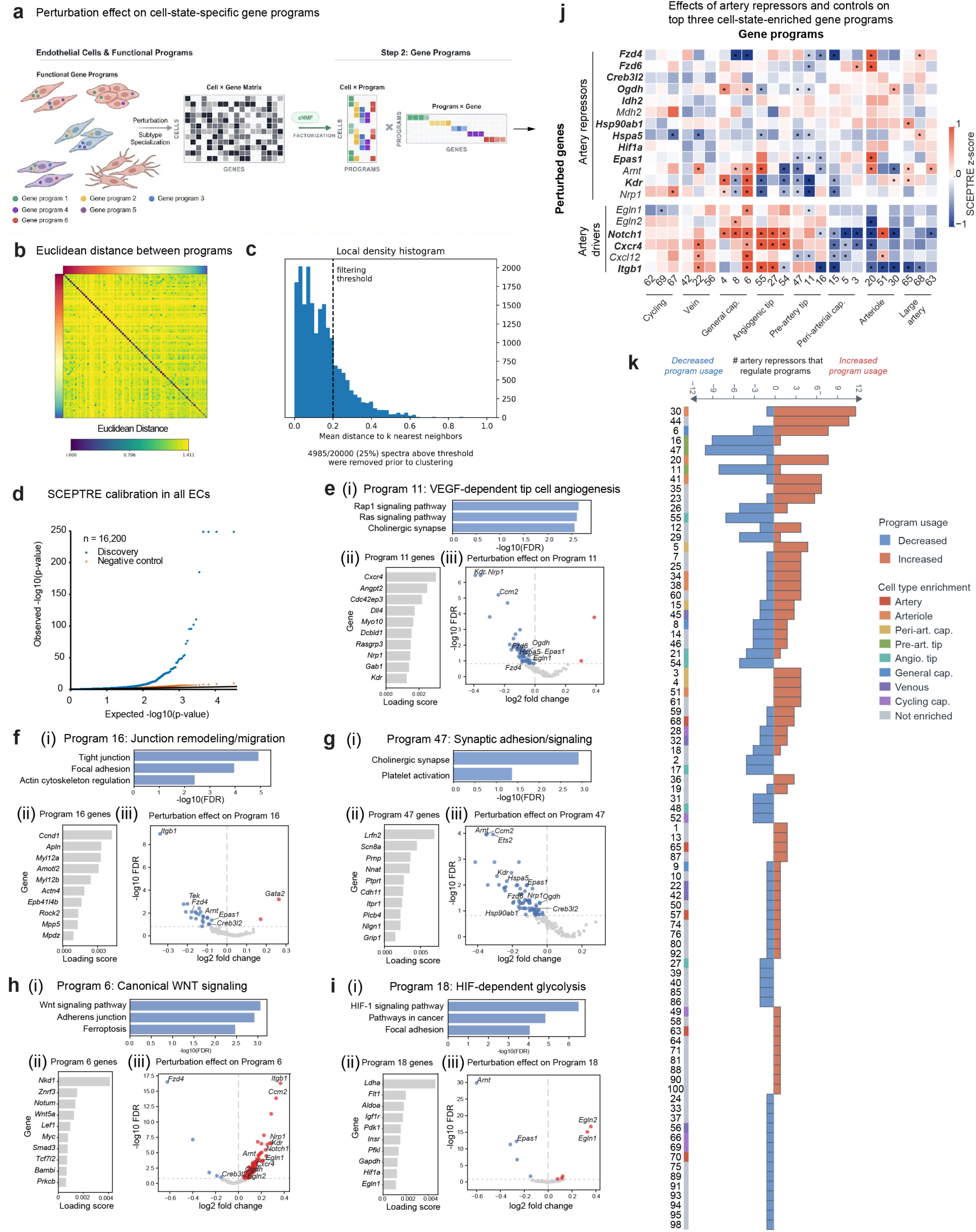
Consensus non-negative matrix factorization (cNMF) identifies cell-subtype-enriched endothelial gene programs regulated by artery repressors. **a**, Schematic of the cNMF workflow used to derive endothelial functional gene programs from the Perturb-seq single-cell expression matrix. A cell × gene matrix was factorized into cell × program and program × gene matrices to define co-regulated gene programs. **b,c**, cNMF quality control and spectrum filtering. **b**, Pairwise Euclidean distances between program spectra across runs. **c**, Distribution of the local density metric (mean distance to k nearest neighbors) used to remove isolated or outlier spectra before consensus clustering; the dashed line marks the filtering threshold. **d**, SCEPTRE calibration for program-level differential testing across all endothelial cells (ECs). Q-Q plot compares observed versus expected −log10(P) values for discovery and matched negative-control tests. **e**–**i**, Additional annotations of selected endothelial gene programs. Each example program is summarized by (i) pathway enrichment, (ii) representative high-loading genes, and (iii) perturbation effects on program usage. **e,** Program 11, a VEGF-dependent tip cell angiogenesis program downstream of *Kdr* and *Nrp1*. **f**, Program 16 tracks a junction remodeling/migration state and is suppressed by *Itgb1.* g, Program 47 captures synaptic adhesion/signaling and is suppressed by *Arnt, Ccm2*, *and Ets2.* h, Program 6 represents canonical WNT signaling and is strongly reduced by *Fzd4* knockdown. **i**, Program 18 captures a HIF-responsive glycolytic state, with reduced usage after *Arnt* or *Epas1* knockdown and increased usage after *Egln1* or *Egln2* knockdown. **j**, Heatmap of perturbation effects for selected artery repressors and perturbed genes on the top three gene programs enriched in each endothelial cell subtype. Rows show selected perturbed genes grouped as artery repressors or artery drivers. Columns show gene programs grouped by the EC state in which they are most enriched. Color denotes SCEPTRE z-score; asterisks indicate significant perturbation effects. FDR < 0.15. **k**, Gene programs ranked by the total number of artery repressors affecting each program. Orange bars indicate the number of artery repressors whose knockdown increased program usage, blue bars indicate the number whose knockdown decreased program usage, and the color strip beside the program labels denotes the EC subtype in which each program is enriched.

**Figure S7.**
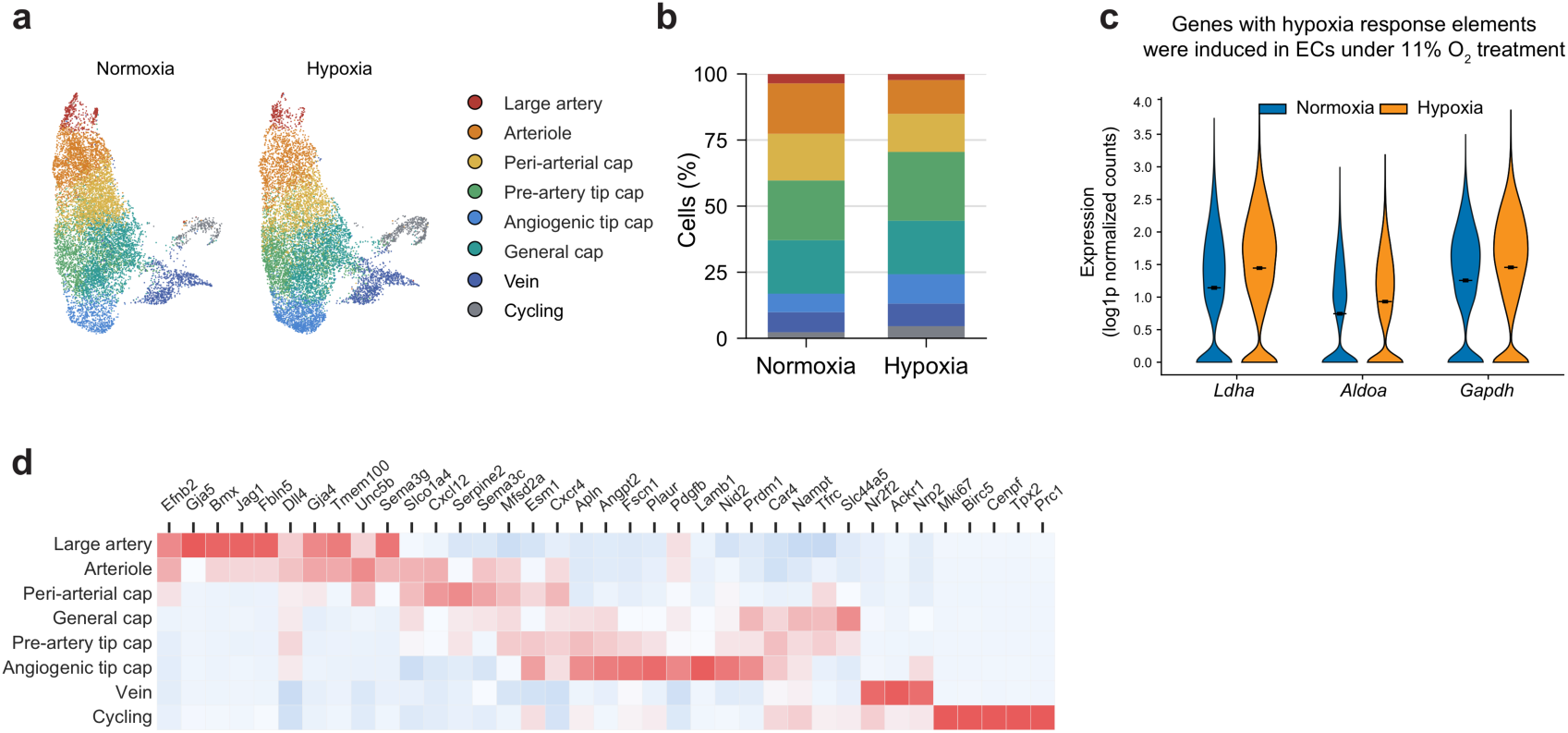
Hypoxia scRNA-seq dataset overview and confirmation of effective neonatal hypoxia exposure. Mouse pups were exposed to normoxia (21% O₂) or mild hypoxia (11% O₂) for 1 week starting at P1, followed by isolation of brain CD31⁺ endothelial cells for scRNA-seq. **a**, UMAPs of endothelial cells from normoxia and hypoxia, colored by annotated endothelial state, showing preservation of the overall endothelial continuum across conditions. **b**, Relative abundance of endothelial states under normoxia and hypoxia. Stacked bars show the fraction of cells in each state; hypoxia shifts the population towards relative expansion of tip-capillary states and lower large artery/arteriole fractions. **c**, Violin plots of *Ldha*, *Aldoa*, and *Gapdh*, representative genes with hypoxia-response elements, showing increased expression under 11% O₂ treatment, consistent with activation of the hypoxia response program. **d,** Heatmap of representative EC subtype marker genes in the hypoxia scRNA-seq dataset.

