## Supplementary material for "A Perturb-seq screen guided by species divergence uncovers pathways for collateral artery formation": Supplementary_material list.docx

**Supplementary Materials**

### Table S1. Marker genes for major cell populations in the integrated embryonic heart scRNA-seq atlas

Generated by Seurat ROC marker testing on the integrated embryonic heart scRNA-seq atlas; this table lists marker genes used to define major cell populations. Columns: cell_type, annotated cell type used for marker calling; gene, marker gene symbol; auc, area under the single-gene ROC classifier; log2fc, log2 fold change for the focal cell type versus all other cells; pct_celltype, fraction of focal cell-type cells expressing the gene; pct_other, fraction of comparison cells expressing the gene; avg_diff, average log-normalized expression difference as reported by Seurat; predictive_power, ROC predictive power calculated as 2 × |AUC − 0.5|; adjusted_p_value, adjusted p-value; n_cells, number of cells in the focal cell type; in_dotplot, whether the gene was included in the representative dot plot.

### Table S2a. Prioritized cross-species endothelial candidate genes for Perturb-seq target nomination

Generated from the cross-species candidate nomination workflow; this table summarizes genes prioritized for Perturb-seq target selection. Tab "S2a Candidate Genes" in Table S2. Columns: gene, gene symbol; nomination_1, first nomination category; nomination_2, second nomination category; nomination_3, third nomination category; selected, inclusion flag for the prioritized export; technical_priority, technical-priority code; full_name, full gene name; heart_p_value, heart endothelial p-value for guinea pig versus mouse; heart_log2fc, heart endothelial log2 fold change for guinea pig versus mouse; heart_pct_gp, percentage of guinea pig heart endothelial cells expressing the gene; heart_pct_mouse, percentage of mouse heart endothelial cells expressing the gene; heart_fdr, heart endothelial FDR; brain_p_value, brain endothelial p-value for guinea pig versus mouse; brain_log2fc, brain endothelial log2 fold change for guinea pig versus mouse; brain_pct_gp, percentage of guinea pig brain endothelial cells expressing the gene; brain_pct_mouse, percentage of mouse brain endothelial cells expressing the gene; brain_fdr, brain endothelial FDR.

### Table S2b. Cross-organ Hallmark pathway enrichment statistics for brain and heart endothelial cells

Generated by Hallmark pathway scoring followed by cross-species Wilcoxon testing in endothelial cells; this table reports pathway-level differences between guinea pig and mouse. Tab "S2b Hallmark Pathways" in Table S2. Columns: pathway, MSigDB Hallmark gene set name; heart_score, Wilcoxon statistic for heart endothelial cells; heart_log2fc, pathway score log2 fold change in heart endothelial cells; heart_p_value, heart endothelial p-value; heart_fdr, heart endothelial Benjamini-Hochberg FDR; brain_score, Wilcoxon statistic for brain endothelial cells; brain_log2fc, pathway score log2 fold change in brain endothelial cells; brain_p_value, brain endothelial p-value; brain_fdr, brain endothelial Benjamini-Hochberg FDR.

### Table S2c. Cross-tissue endothelial differential-expression results supporting Perturb-seq candidate nomination

Generated by matching candidate genes to cross-species differential-expression results from brain and heart endothelial cells; this table indicates whether and how each candidate differs by species in each tissue. Tab "S2c Cross-tissue DEG" in Table S2. Columns: gene, merged gene symbol; gene_key, uppercase token used for matching; brain_gene, matched brain DEG gene symbol; in_brain_deg, whether the gene was present in the brain DEG table; brain_direction, direction of change in guinea pig relative to mouse in brain endothelial cells; brain_score, brain Wilcoxon statistic; brain_log2fc, brain endothelial log2 fold change; brain_p_value, brain p-value; brain_fdr, brain FDR; heart_gene, matched heart DEG gene symbol; in_heart_deg, whether the gene was present in the heart DEG table; heart_direction, direction of change in guinea pig relative to mouse in heart endothelial cells; heart_score, heart Wilcoxon statistic; heart_log2fc, heart endothelial log2 fold change; heart_p_value, heart p-value; heart_fdr, heart FDR.

### Table S2d. Differential endothelial incoming signaling interactions and candidate receptor ranking

Generated from CellChat incoming endothelial interaction analysis combined with ligand and receptor differential-expression statistics; this table ranks ligand-receptor interactions for candidate receptor review. Tab "S2d Signaling" in Table S2. Columns: receptor, endothelial receptor gene; ligand_receptor_pair, ligand-receptor pair; source, sender cell population; target, receiver cell population; ligand, ligand gene; communication_probability, CellChat communication probability; interaction_p_value, CellChat interaction p-value; interaction_id, CellChat interaction identifier; pathway, CellChat pathway name; annotation, CellChat interaction class; evidence, supporting evidence recorded in CellChat; species, species/source dataset label; source_ligand, combined sender-cell and ligand identifier; target_receptor, combined target-cell and receptor identifier; ligand_p_value, ligand differential-expression p-value; ligand_log2fc, ligand log2 fold change for guinea pig versus mouse; ligand_pct_gp, percentage of guinea pig source cells expressing the ligand; ligand_pct_mouse, percentage of mouse source cells expressing the ligand; receptor_p_value, receptor differential-expression p-value; receptor_log2fc, receptor log2 fold change for guinea pig versus mouse; receptor_pct_gp, percentage of guinea pig endothelial target cells expressing the receptor; receptor_pct_mouse, percentage of mouse endothelial target cells expressing the receptor; interaction_log2fc, cross-species interaction log2 fold change; brain_ec_cpm, receptor expression in brain endothelial cells in counts per million; selected, candidate-review flag.

### Table S2e. Cross-species transcription factor ranking for artery repressor candidates

Generated from the cross-species transcription factor prioritization workflow for putative artery repressors; this table compares observed scores with matched random-background scores. Tab "S2e TF Repressors" in Table S2. Columns: row, original row index; gene, transcription factor gene symbol; mouse_score, mouse prioritization score; mouse_background, matched random-background mouse score; mouse_p_value, mouse p-value; mouse_fdr, mouse FDR; gp_score, guinea pig prioritization score; gp_background, matched random-background guinea pig score; gp_p_value, guinea pig p-value; gp_fdr, guinea pig FDR; composite_score, combined cross-species prioritization score; arterial, arterial endothelial expression summary; capillary, capillary endothelial expression summary; venous, venous endothelial expression summary; brain_ec_cpm, expression in brain endothelial cells in counts per million; selected, follow-up selection flag.

### Table S2f. Cross-species transcription factor ranking for artery-essential candidates

Generated from the cross-species transcription factor prioritization workflow for putative artery-essential factors; this table compares observed scores with matched random-background scores. Tab "S2f TF Essential" in Table S2. Columns: row, original row index; gene, transcription factor gene symbol; mouse_score, mouse prioritization score; mouse_background, matched random-background mouse score; mouse_p_value, mouse p-value; mouse_fdr, mouse FDR; gp_score, guinea pig prioritization score; gp_background, matched random-background guinea pig score; gp_p_value, guinea pig p-value; gp_fdr, guinea pig FDR; composite_score, combined cross-species prioritization score; brain_ec_cpm, expression in brain endothelial cells in counts per million; selected, follow-up selection flag.

### Table S3. Custom spatial transcriptomics probe sequences for mouse and guinea pig gene panels

Generated from the custom spatial transcriptomics probe design export; this table provides the gene panel and probe sequences used for mouse and guinea pig spatial validation. Columns: gene, target gene name; probe_sequence, probe oligonucleotide sequence.

### Table S4. List of gRNA sequences in the Perturb-seq library

Generated from the Perturb-seq feature-reference file; this table defines the guide features used for gRNA capture and target assignment. Columns: feature_id, guide or feature identifier; guide_name, guide name; read, sequencing read used for guide capture; pattern, fixed guide-capture pattern adjacent to the barcode placeholder; guide_sequence, captured guide sequence; feature_type, feature class recorded in the feature-reference file; target_id, target gene identifier; target_gene_name, target gene name.

### Table S5. Ranked marker genes for endothelial clusters in the brain vascular Perturb-seq atlas

Generated by ROC marker testing for endothelial clusters in the neonatal brain Perturb-seq dataset; this table lists ranked marker genes for endothelial states. Tab "S5a EC Cluster Markers" in Table S5. Columns: cell_state, endothelial state used for marker selection; gene, marker gene symbol; rank, within-state marker rank; auc, ROC classifier area under the curve; log2fc, log2 fold change for the focal state versus other endothelial states; pct_celltype, fraction of focal-state cells expressing the gene; pct_other, fraction of comparison cells expressing the gene; min_log2fc, log2 fold-change threshold used for marker selection

### Table S6a. Target-level on-target knockdown efficiency and downstream discovery summary for the Perturb-seq screen

Generated by summarizing target-level on-target SCEPTRE results and downstream program-level discoveries from the in vivo CRISPRi Perturb-seq screen; this table reports perturbation QC and discovery yield for each target. Tab "S6a Knockdown Efficiency" in Table S6. Columns: target, perturbed target gene; log2fc, on-target log2 fold change; p_value, on-target p-value; significant, on-target significance call; pct_knockdown, inferred percent knockdown; n_programs, number of program-level rows tested for the target; n_significant_programs, number of significant program-level discoveries.

### Table S6b. Target-specific effects on endothelial UCell state scores in the Perturb-seq screen

Generated by SCEPTRE testing of perturbation effects on endothelial UCell state scores; this table reports target-specific effects on endothelial state signatures. Tab "S6b UCell Scores" in Table S6. Columns: target_order, target ordering index; target, perturbed target gene; response_order, score ordering index; ucell_score, endothelial UCell signature score; log2fc, log2 fold change in perturbed versus control cells; z_score, SCEPTRE z score; p_value, SCEPTRE p-value; fdr, Benjamini-Hochberg FDR within each target across scores; pass_qc, pairwise SCEPTRE QC flag; n_treatment, number of treatment cells with nonzero response values; n_control, number of control cells with nonzero response values.

### Table S7. Pial artery collateral comparison across genotypes

Generated from per-animal pial artery collateral counts in validation comparisons; this table reports the measured collateral counts by comparison group. Columns: panel_id, comparison-panel identifier; panel, perturbation-versus-control comparison label; group, genotype or perturbation group for the animal; collateral_count, counted number of pial collateral arteries.

### Table S8. SCEPTRE results for gRNA-target effects on gene programs in all endothelial cells

Generated by program-level SCEPTRE testing of gRNA-target effects across all endothelial cells; this table reports perturbation effects on cNMF gene programs. Columns: program, cNMF gene program identifier; target, perturbed target gene; n_treatment, number of treatment cells with nonzero program-response counts; n_control, number of control cells with nonzero program-response counts; pass_qc, pairwise SCEPTRE QC flag; p_value, raw SCEPTRE p-value; log2fc, log2 fold change in perturbed versus control cells; stage, SCEPTRE adaptive resampling stage; z_score, observed SCEPTRE z score; xi, skew-normal location parameter for stage-2 tests and NA otherwise; omega, skew-normal scale parameter for stage-2 tests and NA otherwise; alpha, skew-normal shape parameter for stage-2 tests and NA otherwise; fdr, Benjamini-Hochberg FDR within each target across tested programs.

### Table S9a. Annotated summary of 100 Perturb-seq endothelial gene programs

Generated by annotating cNMF-derived endothelial gene programs from the Perturb-seq dataset; this table summarizes the biological interpretation of each program. Tab "S9a Program Summary" in Table S9. Columns: program, cNMF program identifier; program_name, curated program name; top_genes, top loading genes used for interpretation; summary, biological interpretation of the program.

### Table S9b. Ranked loading genes for 100 Perturb-seq gene programs

Generated from cNMF program loading matrices; this table lists the highest-loading genes for each program. Tab "S9b Loading Genes" in Table S9. Columns: gene, gene symbol; score, cNMF loading score for that gene within the program; program, cNMF program identifier.

### Table S9c. Endothelial cell-state enrichment and depletion of Perturb-seq gene programs

Generated by testing cNMF program usage across endothelial states; this table reports state enrichment or depletion of gene programs. Tab "S9c State Enrichment" in Table S9. Columns: cell_state, endothelial state; direction, whether the program is enriched or depleted in that state; program, cNMF program identifier; rank, within-state rank among programs in the same direction; auc, ROC classifier area under the curve; log2fc, log2 fold change of program usage in the focal state versus other states; pct_celltype, fraction of focal-state cells with nonzero program usage; pct_other, fraction of comparison cells with nonzero program usage; jaccard, mean top-gene Jaccard similarity to peer programs selected for the same state and direction.

### Table S10. Differential usage of endothelial gene programs between hypoxia and normoxia

Generated by differential testing of row-normalized cNMF program usage between hypoxia and normoxia endothelial cells using two-sided Mann-Whitney U tests; this table identifies programs altered by neonatal hypoxia exposure. Columns: program, cNMF gene program identifier; mean_hypoxia, mean normalized usage in hypoxia endothelial cells; mean_normoxia, mean normalized usage in normoxia endothelial cells; mean_difference, mean_hypoxia minus mean_normoxia; log2fc, log2 fold change in hypoxia versus normoxia; p_value, Mann-Whitney U p-value; u_statistic, Mann-Whitney U statistic; pct_hypoxia, fraction of hypoxia endothelial cells with usage above 0.01; pct_normoxia, fraction of normoxia endothelial cells with usage above 0.01; max_pct, larger detection fraction across conditions; fdr, Benjamini-Hochberg FDR across all programs.

### Table S11. Differential usage of endothelial gene programs between guinea pig and mouse brain

Generated by differential testing of row-normalized cNMF program usage between guinea pig and mouse brain endothelial cells using two-sided Mann-Whitney U tests; this table identifies gene programs that differ between species in brain endothelium. Columns: program, cNMF gene program identifier; direction, direction of program usage in guinea pig relative to mouse; mean_gp, mean normalized usage in guinea pig brain endothelial cells; mean_mouse, mean normalized usage in mouse brain endothelial cells; log2fc, log2 fold change for guinea pig versus mouse; fdr, Benjamini-Hochberg FDR across all programs.
